# Chemical Compensation to Mechanical Loss in Cell Mechanosensation

**DOI:** 10.1101/2025.04.02.646640

**Authors:** Qin Ni, Zhuoxu Ge, Anindya Sen, Yufei Wu, Jinyu Fu, Alice Amitrano, Nitish Srivastava, Konstantinos Konstantopoulos, Sean X. Sun

## Abstract

Mammalian cells sense and respond to environmental changes using a complex and intelligent system that integrates chemical and mechanical signals. The transduction of mechanical cues into chemical changes modulates cell physiology, allowing a cell to adapt to its microenvironment. Understanding how the chemical and mechanical regulatory modules interact is crucial for elucidating mechanisms of mechanosensation and cellular homeostasis. In this study, we find that cells exhibit non-monotonic changes in cell volume and intracellular pH when subjected to physical stimuli and varying degrees of actomyosin cytoskeleton disruption. We discover that these non-monotonic responses are mediated by a chemical compensation mechanism, where the attenuation of actomyosin activity stimulates the activity of PI3K/Akt pathway. This, in turn, activates sodium-hydrogen exchanger 1 (NHE1), resulting in elevated intracellular pH and increased cell volume. Furthermore, we identify a competitive interaction between the PI3K/Akt and MAPK/ERK pathways - two major regulators of cell proliferation and motility. This competition modulates the chemical compensation based on the relative activities of these pathways. Our mathematical modeling reveals the network structure that is essential for establishing the non-monotonic response. Interestingly, this regulatory system is altered in HT1080 fibrosarcoma, highlighting a potential mechanistic divergence in cancer cells in contrast to their normal-like counterpart, such as NIH 3T3 and HFF-1 fibroblasts. Overall, our work reveals a compensatory mechanism between chemical and mechanical signals, providing a novel infrastructure to elucidate the integrated mechanochemical response to environmental stimuli.

## Introduction

Mammalian cells employ a complex regulatory system that combines chemical and mechanical inputs to modulate cell function and behavior. Cells sense the physical properties of their microenvironment and convert mechanical signals to biochemical activities (*1, 2*). This mechanochemical signaling process leads to changes in cell shape, volume, and properties of the cytoplasm and organelles, enabling cells to adapt to the microenvironment (*3–8*). Revealing the roles of key regulators in this mechanochemical crosstalk is important for understanding mechanosensation and cellular homeostasis.

Studies on mechanosensation over the last two decades have consistently reported that mammalian cells display non-monotonic responses to extracellular mechanical stiffness. Earlier work on cell migration across substrates of varying stiffness (durotaxis) reveals that cell migration speed peaks at an intermediate stiffness (*3, 9, 10*). Non-monotonic changes have also been reported in cell morphology, collective migration, and cytoskeletal mechanics under varying substrate stiffness or other mechanical conditions (*11–20*). More recent work also demonstrated that the impact of substrate stiffness can be biphasic, with stem cell reprogramming efficiency and epigenetic modifications peaking also at an intermediate stiffness (*8*). While models focusing on cell mechanics have been employed to explain the non-monotonic dynamics of cell migration (*10, 21–23*), the recently observed biphasic behavior in epigenetic modifications and reprogramming is likely controlled by a more complex system that integrates cell mechanosensation and chemical signaling cascades.

Among the numerous components involved in integrating mechanical and chemical signals, Na^+^-H^+^ exchanger 1 (NHE1) is likely to be a critical hub. Utilizing the existing Na^+^ gradient across the cell membrane, NHE1 exports protons to regulate intracellular pH. Additionally, NHE1 has been recognized as a key regulator of cell volume and migration due to its role in generating transmembrane water flux (*7, 24–29*), and dysfunction in NHE1-mediated pH and volume regulation is associated with numerous cancers (*7, 30, 31*). NHE1 has been shown to bind to actin filaments at the cell membrane through its co-binding partner ezrin, thereby directly interacting with the actomyosin network (*32, 33*). Our recent work revealed that hypotonic shocks can trigger cytoskeletal activation of NHE1 in normal-like cell lines such as NIH-3T3, leading to an unexpected secondary volume increase (SVI) and alkalinization (*7*). Cell substrate stiffness has also been reported to modulate NHE1-mediated intracellular pH (*34*). On the other hand, NHE1 can be activated directly by Akt, a key component of the PI3K/Akt/mTORC signaling pathway (*35*). These findings suggest that NHE1 is sensitive to both actomyosin activity and the extracellular mechanical environment.

In this work, we reveal a complex, non-monotonic interaction among NHE1, actomyosin network, and PI3K/Akt signaling cascades that governs cell mechanosensation. While mild disruption of the actomyosin network inhibits NHE1 and its corresponding pH and SVI dynamics, as established in our recent work (*7*), severe disruption of the actomyosin network reactivates NHE1, leads to alkalinization and a restoration of SVI under hypotonic shock. Combining mathematical modeling and experiments, we find that this non-monotonic response arises from a compensatory mechanism via PI3K/Akt, a phenomenon we term “chemical compensation to mechanical loss”. Severe actomyosin disruption increases PI3K activity and Akt phosphorylation, which directly activates NHE1 to compensate for the loss of cytoskeletal activation. Our mathematical modeling further reveals the requirements of the actomyosin-PI3K/Akt-NHE1 network necessary to generate chemical compensation in NIH-3T3 cells, and why this response is absent in HT1080 fibrosarcoma cells. We further discover a competition between PI3K/Akt and MAPK/ERK pathways, where the ERK inhibition fine-tunes the activation of the chemical compensation. Overall, we demonstrate how cellular mechanical, biochemical, and electrophysiological elements interact to generate complex dynamics, highlighting the networked nature of a cell systems in response to environmental change.

## Results

### Actomyosin activity regulates NHE1 non-monotonically

We first investigated the role of the actomyosin cytoskeleton in regulating NHE1 by directly disrupting actomyosin using various pharmacological agents and monitoring changes in the intracellular pH (pH_i_). Treatment with a high dose (2 µM) of the actin-depolymerizing agent Latrunculin A (LatA) in NIH 3T3 cells grown on glass rapidly disassembled the actin network within 5 minutes (Fig. 1a) and increased pH_i_ (Fig. 1b). The alkalinization peaked 10 minutes after LatA addition and gradually recovered. To ensure that the observed alkalinization was not specific to LatA, we attenuated cell contractility using the ROCK inhibitor Y-27632 (Y-27). At 20 µM, Y-27 similarly triggered alkalinization (Fig. S1e, f). These results were further confirmed using the pH-sensitive fluorescent dye pHrodo-Red AM (Fig. S1b). To assess whether this alkalinization was mediated by NHE1 activation, we pre-treated 3T3-pHRed cells with 40 µM of the NHE1 inhibitor EIPA prior to LatA (2 µM) or Y-27 treatment. Pre-inhibition of NHE1 completely abolished the alkalinization and, instead, triggered a decrease in pH_i_ (Fig. 1c). We also observed Y-27-induced alkalinization in other cell lines, including hTERT-immortalized retinal pigmented epithelial (RPE-1) cells and human foreskin fibroblast (HFF-1) cells (Fig. 1f). These consistent results, across different pharmacological treatments that target distinct cytoskeletal components, confirmed that NHE1 is sensitive to actomyosin cytoskeleton activity.

**Figure 1.**
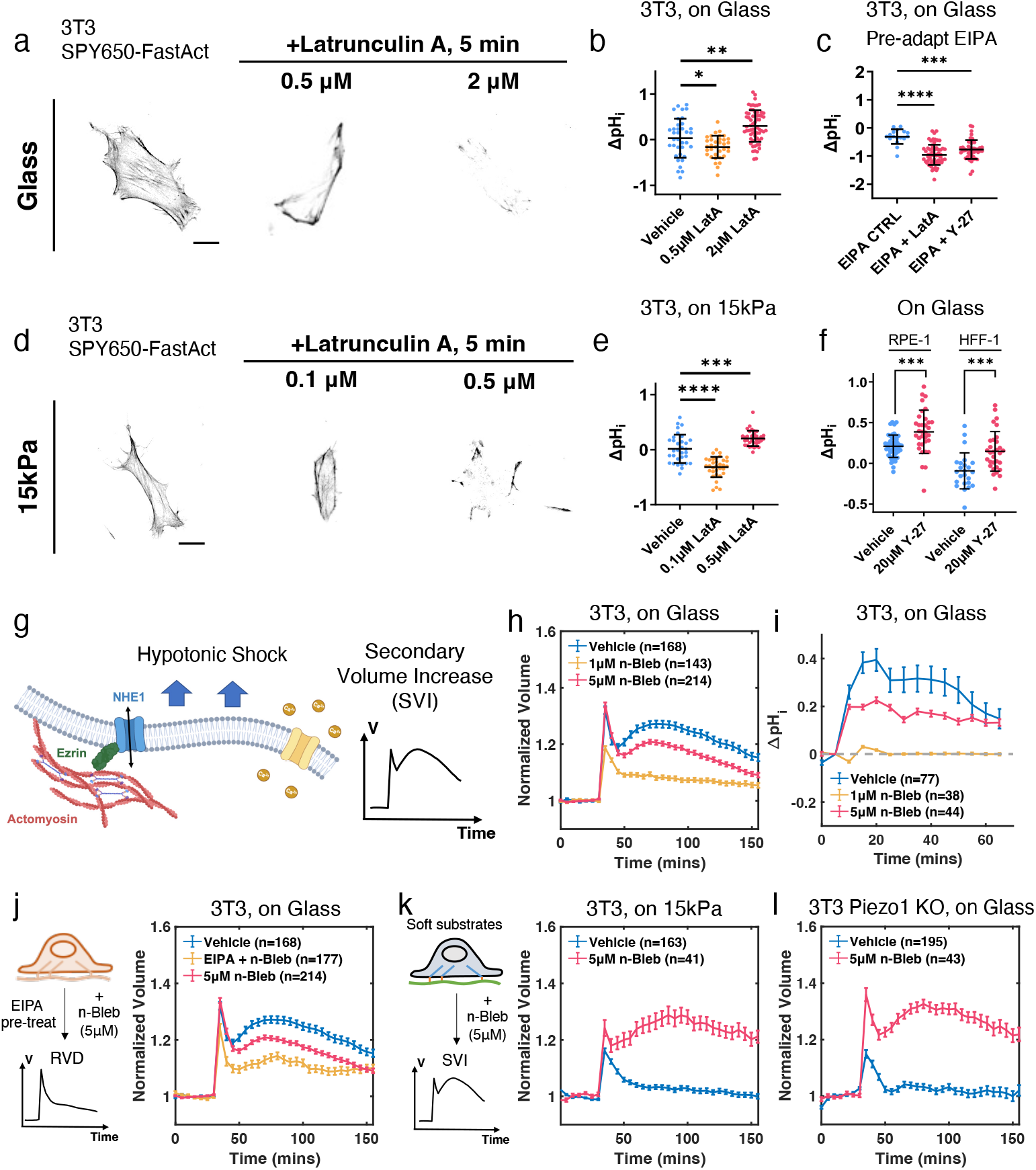
Actomyosin regulates NHE1 activity non-monotonically. (a,d) Representative confocal images of NIH 3T3 on glass (a) and on 15kPa PDMS substrates (d) stained with the live-cell F-actin probe SPY650-FastAct without treatment, and with different dosage Latrunculin A (LatA) treatment for 5 minutes. (b, e) Changes in ΔpH_i_ in 3T3-pHred cells treated with vehicle versus different doses of LatA on glass (b; n = 36, 35, 65) and on 15kPa PDMS substrates (e; n = 32, 33, 35). (c) ΔpH_i_ of 3T3-pHred cells pre-treated with 40 µM EIPA for 2 h prior to vehicle, Y-27, and LatA (2 µM) treatment. n = 17, 65, 48. (f) ΔpH_i_ of RPE-1 and HFF-1 cells treated with vehicle and 20 µM Y-27 for 1 h. n = 49, 34 for RPE-1, and n = 24, 31 for HFF-1. (g) An illustration of the secondary volume increase (SVI) mechanism. Under hypotonic shock, Piezo1 mediated Ca^2+^ influx triggers actomyosin remodeling, leading to NHE1 activation through their binding partner ezrin. (h) Volume dynamics of 3T3 treated with vehicle, 1 µM, or 5 µM n-Blebbistatin (n-Bleb), with hypotonic shock applied at 30 min. All cells were treated 2 h prior to hypotonic shock.(i) ΔpH_i_ of 3T3 treated with vehicle, 1 µM and 5 µM n-Bleb, with hypotonic shock applied at 5 min. (j) Comparison of the 3T3 volume dynamics treated with vehicle, 5 µM n-Bleb, and a combination of 5 µM n-Bleb and 40 µM EIPA, with hypotonic shock applied at 30 min. (k,l)Volume dynamics of 3T3 grown on 15kPa PDMS substrates (g) and 3T3 Piezo1 KO cells (g) treated 5 µM n-Bleb, with hypotonic shock applied at 30 min. (b, c, e, f) Error bars represent STD. Mann Whitney U-tests. (h-l) Error bars represent the standard error of mean (SEM). (a, d) Scale bar = 20 µM.

Interestingly, we found that cytoskeletal regulation of NHE1 follows a non-monotonic pattern. Treating NIH 3T3 cells with a lower dose of LatA (0.5 µM), while sufficient to disrupt actin structures over a longer timescale (Fig. S1a), instead reduced pH_i_ (Fig. 1b and Fig. S1c). On the other hand, 0.5 µM LatA was sufficient to rapidly disassemble the actin network and induce alkalinization in NIH 3T3 cells grown on 15 kPa polydimethylsiloxane (PDMS) substrates (Fig. 1d,e, and Fig. S1a,c). On this compliant substrate, an even lower dose of LatA (0.1 µM), which only partially disrupted the actin network, led to acidosis. Given that a soft substrate can attenuate cytoskeletal force (*4*), these dose-dependent responses suggest that NHE1 is sensitive to both intracellular and extracellular mechanical environments, and its regulation follows a non-monotonic pattern. NHE1 activity is downregulated with mild attenuation of cell mechanics but becomes hyperactivated with severe disruption of cell mechanics.

We then explored how this non-monotonic interaction between actomyosin and NHE1 affected cell response to other types of environmental change. Previously, it was demonstrated that actomyosin can activate NHE1 during hypotonic shock in normal-like cells, including NIH 3T3 (*7*). Using the Fluorescence eXclusion method (FXm) (*36, 37*) to track cell volume at single cell resolution, we have shown that the cytoskeletal activation of NHE1 leads to a secondary volume increase (SVI) following an initial regulatory volume decrease and a rise in pH_i_ post shock (Fig. 1g). Moreover, we have reported that low doses of LatA (0.1 µM), F-actin depolymerizer Cytochalasin D (CytoD, 0.5 µM), and myosin II inhibitor (S)-nitro-blebbistatin (n-Bleb, 1 µM) are able to abolish SVI. Building upon these findings, we used this system as a model to further investigate the actomyosin-NHE interaction. Surprisingly, we discovered that higher doses of LatA (0.5 or 2 µM), CytoD (2 µM), or n-Bleb (5 µM) restored SVI (Fig. 1h and Fig. S1g, h). Furthermore, treating 3T3 cells with 5 µM n-Bleb restored the sustained alkalinization during the SVI (Fig. 1i). Pre-inhibiting NHE1 using EIPA was able to abolish n-Bleb (5 µM) induced SVI, confirming that NHE1 activation is still required in this process (Fig. 1j).

The restoration of SVI under severe actomyosin disruption suggested a new mechanism that operates independently of actomyosin-mediated NHE1 activation. To test this hypothesis, we applied hypotonic shock to cells plated on the 15kPa PDMS substrate with disrupted actomyosin activity. We noted that the 15 kPa substrate had been previously shown to abolish SVI. Similar to our observations on glass, treatment with 5 µM n-Bleb or 0.5 µM LatA restored SVI on the 15 kPa substrate (Fig. 1k and Fig. S1i). Additionally, while SVI normally requires calcium influx through the stretch-activated Piezo1 channel, we found that 5 µM n-Bleb or 0.5 µM LatA were also able to restore SVI in 3T3 Piezo1 CRISPR knockout (KO) cells (Fig. 1l and Fig. S1j).

Together, these results suggest that cells maintain two distinct mechanisms to control pH_i_ and cell volume in a complex, non-monotonic manner. One system is mechanosensitive, activated through a pathway involving Piezo1/Ca^2+^-actomyosin-ezrin-NHE1, as detailed earlier (*7*). The other system becomes active when cytoskeletal forces are significantly attenuated, generating a pH_i_ and volume response similar to that of the mechanosensation-driven system. The latter mechanism appears to be independent of the extracellular mechanical environments and may be triggered by compensatory intracellular biochemical events. We termed this phenomenon “chemical compensation to mechanical loss.”

### Mathematical modeling predicts the network structure for chemical compensation

To probe the potential mechanism of this chemical compensation, we developed a mathematical model to explore the interactions between NHE1, actomyosin, and their corresponding biochemical signaling. We hypothesized that there is a three-component network that includes: 1) a mechanical module mainly comprising the actomyosin cytoskeleton, 2) a biochemical module that interacts with NHE1 and actomyosin, and 3) an ionic module mainly comprising NHE1 (Fig. 2a). In the most general setting, all 3 modules can mutually interact with each other. Taking inspiration from neural network models, we describe these modules in a coarse-grained manner using activation functions. The activation of any one module by the other two modules are modeled by the Hill function with an activation threshold. This type activation function is also widely used in biological networks (*38, 39*). The dynamics of module *i* is described by

**Figure 2.**
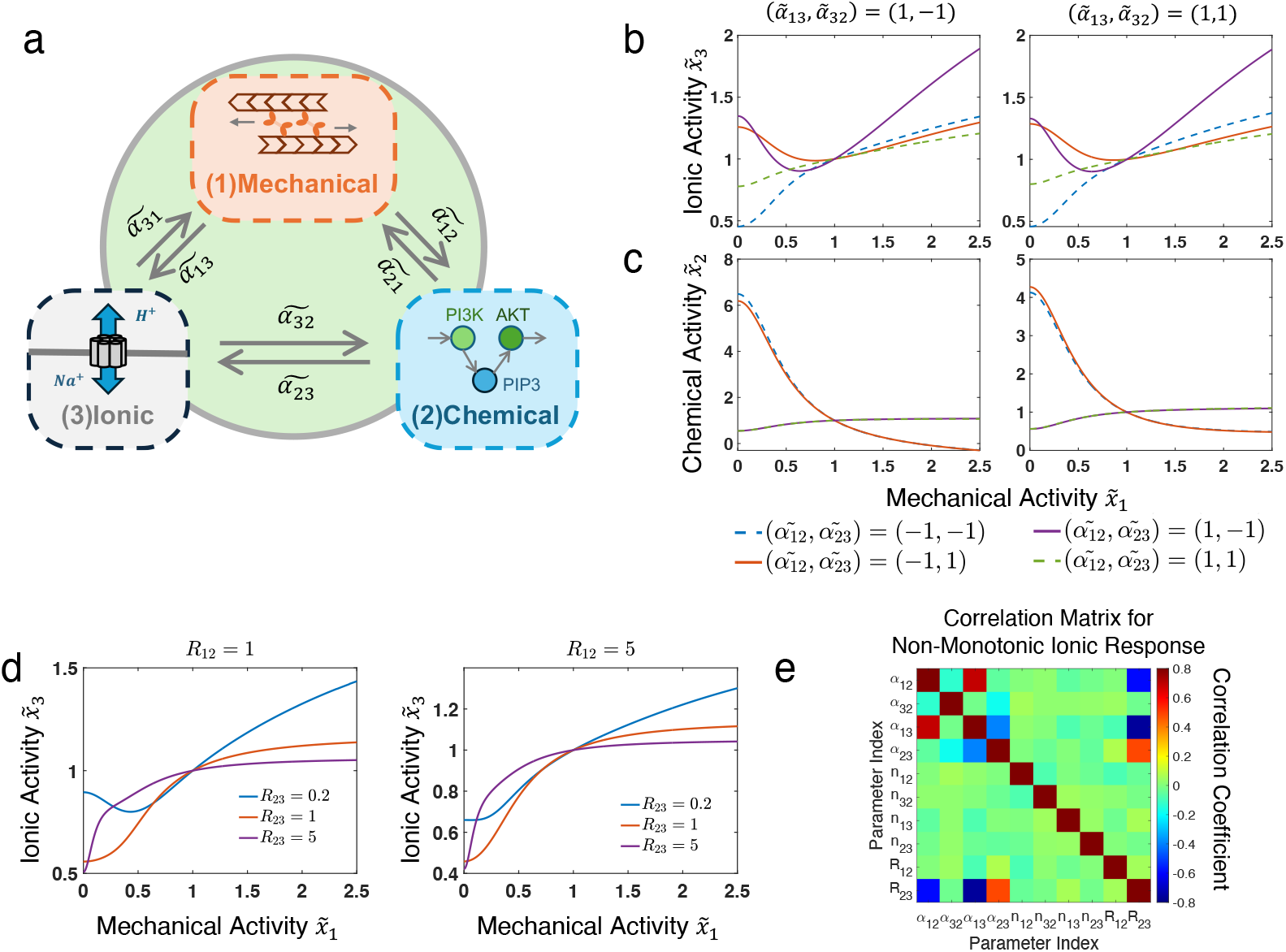
Mathematical modeling reveals the network structure for chemical compensation to mechanical loss. (a) An illustration of the three-component mechanochemical model. (b, c) Activities of the ionic module (b) and the chemical module (c) influenced by mechanical activity 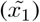 under different net-work structures. *α*_*i,j*_ > 0 indicates module i activates module j, and vice versa. (d) Ionic module activity influenced by mechanical activity at different scaling factors *R*_12_ = *k*_31_*/k*_32_ and *R*_23_ = *k*_12_*/k*_13_. The scaling factor in mechanical-chemical interaction (*R*_23_) should be lower than that in mechanical-ionic interaction (*R*_12_) to generate the non-monotonic behavior. (e) Correlation matrix showing Pearson correlation coefficients for all parameters generating non-monotonic ionic activity from the unbiased parameter sampling (detailed in Supplemental Text). (b-d) If not specified, the base values of all parameters are set as: 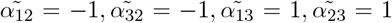, *n*_12_ = *n*_32_ = *n*_13_ = *n*_23_ = ™2, *R*_12_ = *R*_23_ = *R*_31_ = 1*/*5. All results are normalized to a base activity level at 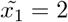

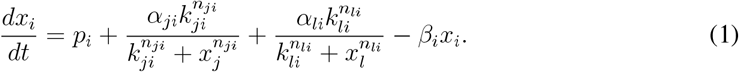

Here, *x*_*i*_ denotes the activity of the *i*-th module. *p*_*i*_ and *β*_*i*_ are the basal activation and deactivation rates, respectively. *α*_*ji*_, *k*_*ji*_, and *n*_*ji*_ describe the activation of *i*-th module by the *j*-th module, which are the magnitude of interaction, scaling factor of activity, and sensitivity of the response, respectively. By assuming that *n*_*ji*_ is always negative, the activation/inhibition process can be solely determined by the sign of *α*_*ji*_. For example, when *α*_*ji*_ < 0, module *j* inhibits the activity of module *i*. Eq. 1 can then be non-dimensionalized as:

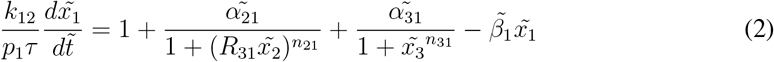

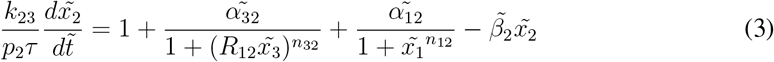

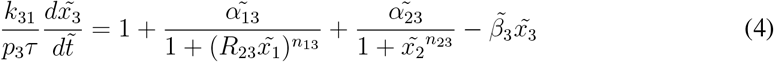

where 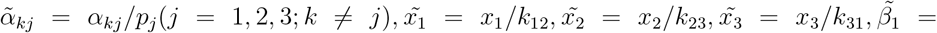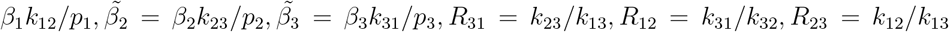 is the time scale of our system. The steady state solution can be obtained by setting all time derivatives to zero. By manipulating one module, we can explore how it influences other two modules and therefore infer the underlying network dynamics. Since we hypothesized that there existed a chemical compensation to mechanical loss, we make actomyosin activity (*x*_1_) as the control variable and explore how it influences the chemical module and ionic activity. Due to the inherent symmetry in the governing equations (Eq.2 - 4), the findings obtained when setting the mechanical module as control variable can be generalized to scenarios where different control variables are employed.

We first explored the ionic module (NHE1) activity as a function of the mechanical (actomyosin) module for different network parameters. To simplify the analysis, we constrained *α*_13_ > 0, reflecting the known activation of NHE1 by actomyosin via ezrin, as previously reported (*7, 32, 33*). Since we selected actomyosin, *x*_1_, as the control variable, we ignored the feed-back to *x*_1_ by setting *α*_31_ = *α*_21_ = 0. Different interactions between ionic-chemical modules (*α*_23_ and *α*_32_), and between mechanical-chemical modules (*α*_12_) were examined, and their resulting activities of the ionic and chemical modules are shown in Fig. 2b and c. In different parameter regimes, monotonic versus non-monotonic ionic responses were observed. We found that, regardless of the ionic module’s influence on the chemical module (*α*_32_), achieving a non-monotonic response requires the net activity of mechanical-chemical-ionic modules to be negative. In other words, interactions between mechanical-chemical modules (*α*_12_) and chemical-ionic modules (*α*_23_) must have opposite signs (*α*_12_ × *α*_23_ < 0; Fig. 2b). In this network, ionic activity first decreases as actomyosin activity decreases, but is re-activated when the mechanical activity falls below a threshold. The magnitude of this non-monotonic change depends on the relative scaling factors of activity *R*_*ji*_ (discussed below). When the interactions between mechanical-chemical modules and ionic-mechanical modules were both positive or both negative, a monotonic ionic response to actomyosin activity was observed. On the other hand, the chemical module activity depends on the feedback from both mechanical and ionic module (*α*_12_ and *α*_32_), but not on its regulation of the ionic modules (*α*_23_; Fig. 2c). We further relaxed the constraint to allow *α*_13_ < 0 to examine scenarios where actomyosin directly inhibits NHE1. In this case, the requirement to achieve non-monotonic ionic activity was reversed, where the net activity of mechanical-chemical-ionic modules must be positive (i.e. *α*_12_ × *α*_23_ > 0; Fig. S2a,b). However, the modeled ionic activity contradicted our experimental observations, and thus we excluded this condition from consideration.

We also examined how variations in the activity scaling factors, *R*_*ji*_, influence ionic activity. *R*_*ji*_ represents both the sensitivity of a module to its input and the magnitude of its response. Our analysis revealed distinct response patterns across different scaling parameter values. Specifically, achieving a non-monotonic compensation response requires *R*_12_ ≤ 1 and *R*_23_ ≤ 1. This observation indicated that actomyosin must influence the chemical module and NHE1 at different scales to achieve effective chemical compensation (Fig. 2d). Similarly, the ionic module must influence the other two components on distinct scales.

We further investigated how chemical and ionic activities depend on all model parameters. Using large-scale unbiased parameter sampling, we calculated the steady-state solution of ionic activity (*x*_3_) for different combinations of parameters. We then defined the ionic module sensitivity as the difference between the maximum and minimum values of ionic activity when varying the mechanical activity *x*_1_, and calculated its correlation with all parameters. We found that the most important parameters influencing ionic sensitivity are *α*_13_, *α*_23_, and *R*_23_ (Fig. S2c). On the other hand, *α*_12_ and *α*_32_ are the most critical parameters influencing the sensitivity of the chemical module (Fig. S2d). Moreover, we found that some pairs of these key parameters display strong correlations with each other (Fig. 2e), revealing an intrinsic relationship between control functions of different modules that are required to regulate the output dynamics in this three-component network. Details of the unbiased parameter sampling and more analysis can be found in the Supplemental Text.

Taken together, the mathematical model predicted the minimal network requirements to achieve chemical compensation during mechanical loss. First, the net influence of actomyosin on NHE1 via the chemical module should be negative (*α*_12_ × *α*_23_ < 0). Second, actomyosin should impact the chemical module and NHE1 at different scales (*R*_23_ ≤ 1). Similarly, NHE1 should impact the other two components at different scales (*R*_12_ ≤ 1). These insights provide a strong rationale for experimentally identifying the chemical module and its interactions with other modules.

### Chemical compensation is achieved through PI3K/Akt activation

From the model predictions, PI3K/Akt emerged as a promising candidate as the chemical modulator within the compensatory network. The PI3K/Akt pathway has a well-established role in regulating actin polymerization (*40, 41*), and NHE1 has been identified as a direct substrate of Akt (*35*). To validate the influence of PI3K/Akt on NHE1, we inhibited PI3K using LY294002 (LY29). Direct treatment with LY29 induced acidosis (Fig. 3d), while pre-treating with LY29 was sufficient to block both Y-27 induced alkalinization and n-Bleb induced SVI (Fig. 3e). Taken together, previous research and our observations suggest a connection between PI3K/Akt, actomyosin, and NHE1, indicating that PI3K/Akt directly activates NHE1 (i.e., *α*_23_ > 0). Under this circumstance, the model predicts that PI3K/Akt must be negatively regulated by actomyosin to satisfy the net negative control (i.e. *α*_12_ · *α*_23_ < 0).

**Figure 3.**
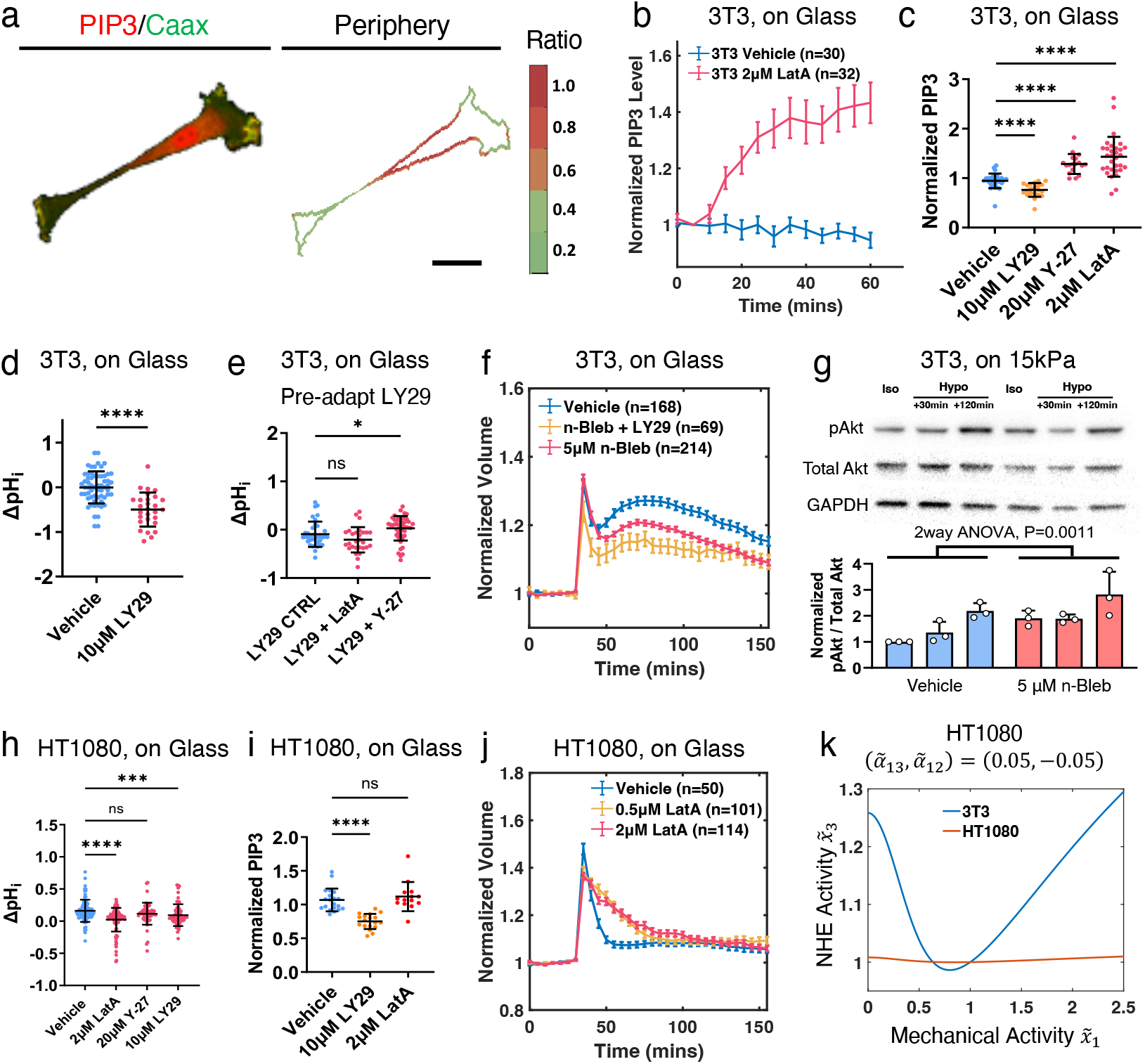
PI3K/Akt chemically compensates the mechanical loss after cytoskeleton disruption. (a) Representative images of 3T3 cells co-transfected with PIP3 indicator (AktPH-mCherry) and membrane indicator (GFP-CaaX), and their ratio at the cell boundary. The ratio at the cell boundary indicates the relative PIP3 level. (b) Normalized PIP3 dynamics of 3T3 cells treated with vehicle control and 2 µM LatA. (c) Normalized PIP3 level of 3T3 treated with vehicle and 2 µM LatA. n = 30, 21, 18, 31. (d) ΔpH_i_ of 3T3-pHred cells treated with vehicle and 10 µM LY294002 (LY29) for 1 h. n = 36, 28. (e) ΔpH_i_ of 3T3-pHred cells pre-treated with 10 µM LY29 for 2 h prior to treatment of vehicle, 2 µM LatA, and 20 µM Y-27. n = 39, 28, 47. (f) Comparison of the 3T3 volume dynamics treated with vehicle, 5 µM n-Bleb, and a combination of 5 µM n-Bleb and 10 µM LY29, with hypotonic shock applied at 30 min. (g) Representative western blot images and quantification of Akt activity in 3T3 cells grown on 15 kPa PDMS substrates treated with vehicle or 5 µM n-Bleb, in isotonic media, and 30 min or 120 min post-hypotonic shock. N = 3 repeats. (h) ΔpH_i_ of HT1080 cells after treating with vehicle control, 2 µM LatA, 20 µM Y-27, and 10 µM LY29 using pHrodo Red-AM. n = 86, 109, 59, 63. (i) Normalized PIP3 level of HT1080 treated with vehicle, 10 µM LY29, and 2 µM LatA. n = 20, 14, 18. (j) Volume dynamics of HT1080 treated with 0.5 and 2 µM LatA. Hypotonic shock was applied at 30 min. All cells were treated 1 h prior to hypotonic shock. (k) NHE1 activity of 3T3 versus HT1080 predicted by the model. 3T3 parameters are set as indicated in Fig. 2. HT1080 parameters are the same as 3T3 excepted for 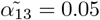 and 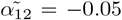 as indicated in the figure. (c-e, g-i) Error bars represent STD. (b, f, j) Error bars represent SEM. (b-e, h-i) Mann Whitney U-tests. (g) Two-way ANOVA test. (a) Scale bar = 20 µM.

Next, we verified the negative controls between actomyosin and PI3K/Akt predicted by the model. To assess PI3K activity, we co-transfected 3T3 cells with the PIP3 indicator AktPH-mCherry and membrane indicator CAAX-GFP. The AktPH to CAAX signal ratio at the cell periphery indicates the membrane bound PIP3 level (Fig. 3a). When LY29 is used to inhibit PI3K, the conversion of PIP2 to PIP3 is blocked, resulting in reduced membrane-bound PIP3 (Fig. 3c and Fig. S3a). In agreement with our prediction, PIP3 levels increased under LatA (2 µM) and Y-27 (20 µM) treatments, indicating that PI3K was hyper-activated (Fig. 3b,c and Fig. S3b). Furthermore, western blot analysis of Akt activity confirmed that both LatA and Y-27 treatments elevated Akt phosphorylation 30 minutes post-treatment (Fig. S3c). Under hypotonic shock, we also found that n-Bleb (5 µM) treatment increased phosphorylated Akt level (pAkt) at the onset of SVI (30 minutes after hypotonic shock) on 15 kPa substrates (Fig. 3g). Pre-inhibiting PI3K/Akt using LY29 reduced n-Bleb induced SVI on glass (Fig. 3f), further supporting the involvement of PI3K/Akt in the chemically compensatory SVI generation. Collectively, these results confirm the presence of a net negative control of actomyosin on NHE1 via PI3K/Akt and its regulation of pH_i_ and SVI, consistent with model predictions.

### HT1080 is insensitive to mechanical loss

While we observed chemical compensation in normal-like cell lines such as 3T3, RPE-1, and HFF-1, this phenomenon was absent in the HT1080 fibrosarcoma cell line. Disruption of actomyosin in HT1080 either reduced pH_i_ (using 2 µM LatA) or had no effect (using 20 µM Y-27; Fig. 3h). Additionally, HT1080 and many other cancer cell lines naturally lack SVI under hypotonic shock (*7*). Even treated with 0.5 µM or 2 µM LatA, no SVI was observed in HT1080 (Fig. 3j). We also observed that while adding LY29 still reduced HT1080 pH_i_ as found in 3T3 (Fig. 3h), LatA failed to generate noticeable changes in membrane PIP3 levels (Fig. 3i). Moreover, western blot showed that NHE1 inhibition using EIPA significantly reduced Akt activity in HT1080, in contrast to the increase seen in 3T3 (Fig. S3d). These striking differences indicated that HT1080 employed a different, actomyosin-insensitive regulatory system.

Our three-component network model provides potential explanations of how HT1080 differs from 3T3. According to the unbiased parameter sampling, *α*_13_ is most critical to the NHE1 sensitivity while *α*_12_ is most critical to PI3K/Akt sensitivity under mechanical loss (Fig. 2e and Fig. S2c). Reducing these two parameters without changing others indeed reproduces the experimental observations of insensitive NHE1 and PIP3 activity in HT1080 under actomyosin disruption (Fig. 3k and Fig. S3e). Interestingly, while reducing *α*_12_ alone is sufficient to abolish the PI3K/Akt response to mechanical loss, insensitive NHE1 response can only be achieved when reducing both *α*_13_ and *α*_12_ simultaneously (Fig. S3f, g).

### ERK/MAPK inhibition stimulates the PI3k/Akt compensation

Lastly, we asked whether the chemical compensation could be activated by stimuli other than mechanical loss. The ERK/MAPK pathway is a major signaling cascade responsible for sensing environmental signals and controlling cellular function. It has been found to exhibit cross-inhibition with the PI3K/Akt pathway. For instance, inhibition of MEK (the upstream regulator of ERK) can enhance growth factor-induced Akt activation (*42, 43*). Utilizing this feature, we inhibited ERK to elevate PI3K/Akt activity and investigate its role in chemical compensation. Similar to the effects of n-Bleb, treatment with 1 µM MEK inhibitor Trametinib (Trame) significantly promoted Akt phosphorylation on soft substrates, as observed by western blot analysis (Fig. 4a). Additionally, Akt activation was further enhanced by Trame 30 minutes after hypotonic shock, resulting in restored SVI and associated alkalinization (Fig. 4b, c). Trame, like n-Bleb, also triggered SVI in 3T3 Piezo1 KO cells (Fig. S3h). To further elucidate the competition between the ERK/MAPK and PI3K/Akt pathways, we treated cells with a combination of Trame and LY29 at varying concentrations on 3T3 grown on the 15kPa PDMS substrate (Fig. 4d). When cells were co-treated with 0.2 µM Trame and 10 µM LY29, SVI was abolished. However, increasing the dosage of Trame to 1 µM overrode the inhibitory effect of LY29, restoring SVI. These results underscore the competing roles of the PI3K/Akt and ERK/MAPK pathways, the two major cell regulatory pathways, in regulating cellular physiological and mechanical responses. The involvement of ERK/MAPK pathways also indicates that the complete mechanochemical regulation of NHE1 is embedded in a complex network that is substantially beyond the simple three-component model discussed here.

**Figure 4.**
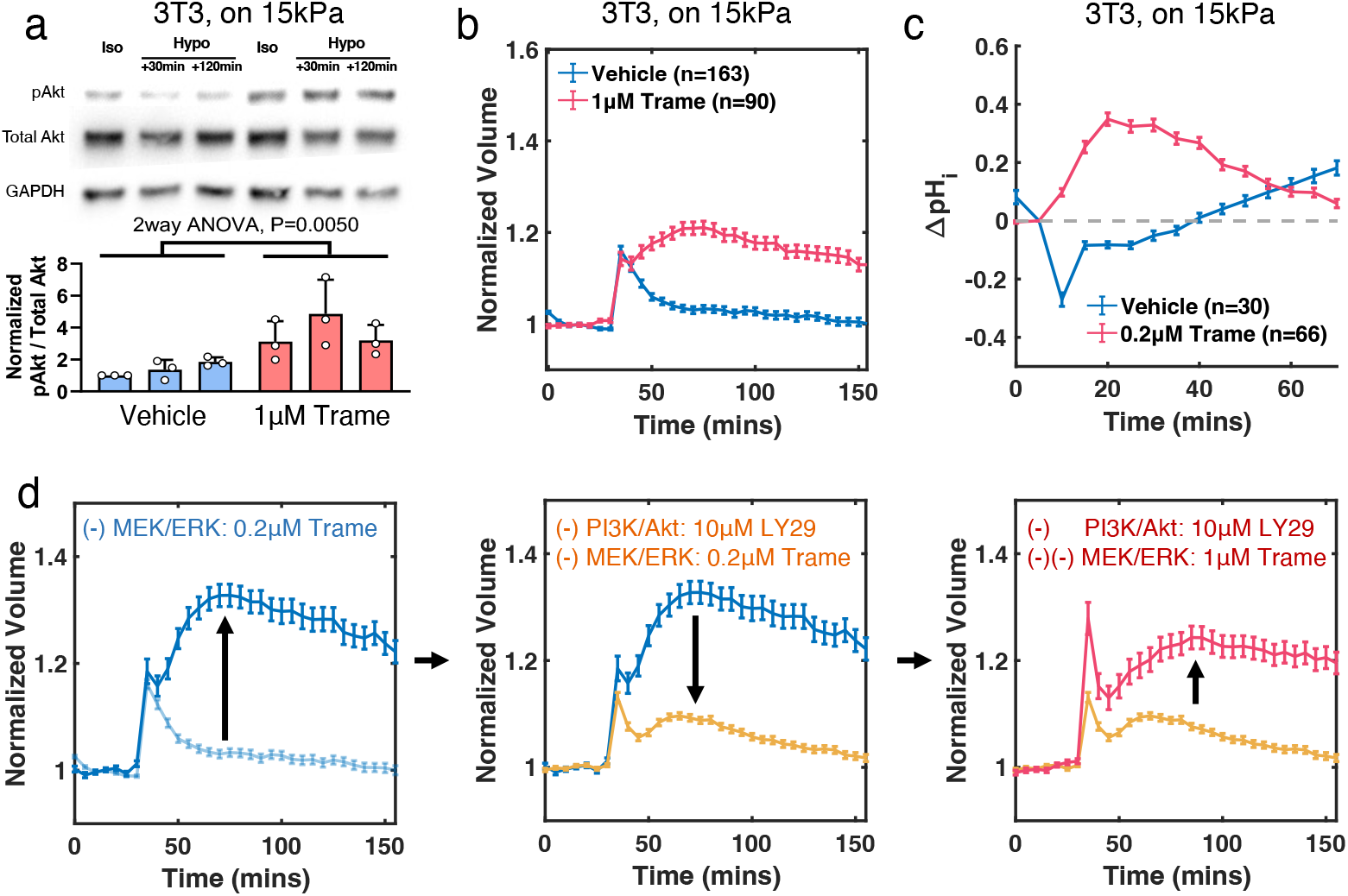
ERK/MAPK inhibition stimulates the PI3k/Akt compensation. (a) Representative western blot images and quantifications of Akt activity in 3T3 cells grown on 15 kPa PDMS substrates treated with vehicle or 1 µM Trametinib (Trame), in isotonic media, and 30 min or 120 min post-hypotonic shock. N = 3 repeats. (b) Volume dynamics of 3T3 grown on 15kPa substrates pre-treated with 1µM Trame. Hypotonic shock was applied at 30 min. (c) ΔpH_i_ of 3T3 grown on 15kPa substrate pre-treated with 1µM Trame using pHrodo Red-AM. Hypotonic shock was applied at 5 min. (d) Comparison of cell volume dynamics under different combination of PI3K/Akt inhibition and MEK/ERK inhibition, with hypotonic shock applied at 30 min. (−)PI3K/Akt = 10 µM LY29, n = 85; (−)MEK/ERK = 0.2 µM Trame, n = 157; and (−)(−)MEK/ERK = 1 µM Trame, n = 64. (a) Error bars represent STD. Two-way ANOVA. (b-d) Error bars represent SEM.

## Discussion

In this work, we demonstrate how a minimal, three-component mechanochemical system interacts to produce a non-monotonic response to environmental stimuli. In this system, NHE1, a central cytoplasmic pH and volume regulator, is simultaneously influenced by a mechanical module (actomyosin) and a chemical module (PI3K/Akt). There are also mutual feedback interactions between all the modules, where we uncovered an important chemical compensation mechanism after mechanical loss. When the integrity of the actin network or myosin contractility is compromised, PI3K/Akt becomes hyper-activated, resulting in NHE1 activation and the corresponding alkalinization. Under hypotonic shock, this chemical compensation also can trigger SVI, a response that is associated with nucleus deformation, epigenetic and transcriptomic modification, and proliferation inhibition (*7*). Interestingly, the PI3K/Akt compensation can also be activated by MAPK/ERK inhibition. This suggests that our three-component mechanochemical regulation of NHE1 is a minimal model embedded in a larger regulatory network that should be examined systematically in the future. Given that PI3K/Akt and MAPK/ERK are two regulators of major signaling pathways responsible for cell proliferation, stress response, and metabolic activity (*43*), it is likely that the larger regulatory network integrates multiple mechanical, chemical, and physiological signals to ultimately regulate all essential cell functions (*31*).

However, have we fully resolved the complex cellular mechanical-chemical-physiological interactions and explained the consequential non-monotonic dynamics? Not entirely. In this work, we focus on NHE1 due to its known interactions with the cytoskeleton and its critical role in cell migration. It is likely that other ion transporters also exhibit similarly complex mechanochemical responses, such as the interaction between K^+^ channels and PIP2/3 (*44*). Moreover, the entire system forms a sophisticated, highly interconnected network, where any observations are likely the result of cooperative interactions within this system. We are probably far from understanding the full picture of this system. For example, the precise mechanisms underlying chemical compensation remain unclear, and it is not yet understood the molecular details of why certain cell types, such as HT1080, lack this compensatory response. The multi-component networked nature of this system makes it inherently unpredictable, because any omission of a seemly unrelated component could significantly alter the network behavior. How, then, can we unravel this mystery without fully knowing its scope? To move forward, a substantial change in research infrastructure is needed, where different experiments generated by different facets of the system should be combined in an unbiased manner (*45*).

## Materials and Methods

### Cell Culture

NIH 3T3, HT1080, and HFF-1 were purchased from the American Type Culture Collection (ATCC). RPE-1 cells were a gift from Rong Li (Johns Hopkins University, Baltimore, MD). All cells were cultured in Dulbecco’s modified Eagle’s media (Corning) supplemented with 10% fetal bovine serum (FBS; Sigma), and 1% antibiotics solution containing 10,000 units/mL penicillin and 10,000 µg/mL streptomycin (Gibco). All cell cultures and live cell experiments were conducted at 37 ^°^C and 5% CO2.

### Osmotic shock media preparation and pharmacological inhibitors

Before the hypotonic shock experiment, cells were pre-incubated in an isotonic solution overnight. The isotonic solution (312 mOsm) contains 50% Dulbecco’s phosphate-buffered saline without calcium and magnesium (Sigma-Aldrich) and 50% cell culture media. 50% hypotonic media (175 mOsm) was prepared by mixing 50% ultra-pure water with 50% cell culture media. The osmolality was measured using Advanced Instruments model 3320 osmometer.

In select experiments, cells were treated with the following pharmacological agents: DMSO as vehicle control (0.1%, Invitrogen), Y-27632 (Tocris), Latrunculin A (Tocris), Cytochalasin D (Tocris), (S)-nitro-Blebbistatin (Cayman Chemical), EIPA (Tocris), LY294002 (Sigma), and Trametinib (Medchemexpress). The doses of inhibitors were stated in the main text. In osmotic shock experiments, cells were pre-adapted to pharmacological agents for 2h except for Latrunculin A and Cytochalasin D (1 h).

### Microfluidic device fabrication

A detailed protocol can be found in Ref. 37. In brief, FXm channel masks were designed using AutoCAD and ordered from FineLineImaging. Silicon molds were fabricated using SU8-3010 (Kayaku) photoresist following standard photolithography procedures and manufacturer’s protocol. Two layers of photoresist were spin coated on a silicon wafer (IWS) at 500 rpm for 7 s with an acceleration of 100 rpm/s, and at 2,000 rpm for 30 s with acceleration of 300 rpm/s, respectively. After a 4 min soft bake at 95 ^°^C, UV light was used to etch the desired patterns from the negative photoresist to yield feature heights that were ∼12 µm. The length of the channels is 16 mm and the width is 1.2 mm.

A 10:1 ratio of PDMS Sylgard 184 silicone elastomer and curing agent were vigorously stirred, vacuum degassed, poured onto each silicon wafer, and cured in an oven at 80 ^°^C for 45 min. Razor blades were then used to cut the devices into the proper dimensions, and inlet and outlet ports were punched using a blunt-tipped 21-gauge needle (McMaster Carr). The devices were cleaned by sonicating in 100% isopropyl alcohol for 10 min, and dried using a compressed air gun. The devices and sterilized 50-mm glass-bottom Petri dishes (FlouroDish Cell Culture Dish; World Precision Instruments) were exposed to oxygen plasma for 1 min for bonding. The bonded devices were then placed in an oven at 80 ^°^C for 45 min to further ensure bonding.

### Polydimethylsiloxane (PDMS) substrates fabrication

To generate 15 kPa PDMS substrates, 2% Sylgard 184 (Dow, 10:1 base to curing agent ratio) were added to Sylgard 527 (Dow, 1:1 base to curing agent ratio) (*46*). In all cases, the elastomer was vacuum-degassed for approximately 5 min to eliminate bubbles, and then spin-coated onto 35 mm or 50 mm glass bottom dishes at 1,000 rpm for 60 s. The dishes were cured 24 h at room temperature. The Young’s modulus was confirmed using a rheometer. Some devices were subsequently plasma-treated and bonded to the FXm devices for cell volume measurement. Before seeding cells, all devices were sterilized using UV for at least 15min.

### Cell volume tracking experiment

Micro-fluidic Fluorescence Exclusion method (FXm) chambers were incubated with 50 µg/mL of type I rat-tail collagen (Enzo) for 1 h at 37 ^°^C, followed by washing with PBS. Before experiments, ∼ 1-2 million per mL cells with 0.2 mg/mL Alexa Fluor 488 Dextran (MW 2,000 kDa; ThermoFisher) dissolved in isotonic media were injected into the devices using a syringe. Cells were then incubated for 1-2 h to allow them to attach and spread. To apply osmotic shock, media were injected into the channel gently using a syringe with the same amount of dextran. For the osmotic shock experiment with drug treatment, cells were pre-adapted 1-4 h in the FXm device before applying osmotic shock. Cells were imaged using fluorescence microscopes as detailed below. The microscope was also equipped with a CO2 module and TempModule stage top incubator (Pecon) that was set to 37 ^°^C and 5% CO2 during the experiment.

### Single cell tracking, cell volume calculation, and image analysis algorithms

Individual cells were tracked by customized MATLAB code using the following algorithm. Firstly, a rough cell mask was drawn from a Gaussian blurred fluorescence image by a set threshold intensity for each cell. The cell mask was then expanded by 10 pixels (2.27 µm) in each direction to ensure proper cell cropping, and a rectangular box was drawn based on the largest lateral dimensions in the x and y directions. A cell would be discarded if any overlapping cells were found in the box. After all images at different time points were processed, we ran an automatic cell tracking algorithm based on the position of the geometric center of each cell mask. In brief, the algorithm assigned each cell (*C*_*t*_) to the closest cell in the next time frame (*C*_*t*+Δ*t*_), and then cross validated the tracking by assigning *C*_*t*+Δ*t*_ to its closest cell at frame t. The cell would be discarded if the cross validation failed to find *C*_*t*_ as the closest neighbor or found more than one closest neighbor, which is normally due to cell migration, detachment, or overlapping with other cells. The results from the automatic cell tracking algorithm were then examined manually by one of the authors.

Cropped images from single cell tracking were then used for cell volume calculation. The mean fluorescence intensity of the pixels outside cell masks, defined as the mean background intensity *I*_*bg*_, reflects the height of the FXm channel. The local intensity within the cell mask, *I*_*V*_ defined the difference between channel height and cell height. Given a known channel height *h*, measured by confocal microscopy, the cell volume *V* can be calculated using the equation 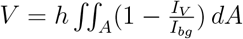. The accuracy of FXm has been tested in our previous work (*7*).

We used a consistent approach for cropping and background subtraction to ensure accuracy and reproducibility. For both confocal or epi-fluorescence images, a rectangle was cropped for each cell, and the boundary of the cropped area was used for calculating background intensity for each channel to reduce noise and enhance signal clarity. To assess the PIP3 level, a cell mask was generated by thresholding, and the cell boundary was determined as the outermost layer of the mask. The average intensities of PIP3 and membrane indicators within the boundary mask were calculated, and the ratio was used to determine the PIP3 level. For other experiments, a cell mask was generated by thresholding, and the average intensity within the mask was calculated.

### Intracellular pH measurement

In this work, we used two methods to assess intracellular pH. In experiments using the ratiometric indicator pHred, cell lines stably expressing pHred were generated as described below. pHred cells were excited at 405 and 561 nm, and emissions above 550 nm were collected using a Zeiss LSM800 confocal microscope. Intracellular pH is inversely proportional to the intensity ratio at 561 over 405 nm. In selected experiments, intracellular pH was measured using pHrodo Red AM (Invitrogen), following the manufacturer’s instructions. In brief, cells were then incubated with media containing a dilution of 5 mM pHrodo Red AM and 1:100 dilution of PowerLoad (100X; Invitrogen) at 37 °C for 30 min. The cells were then gently washed once with media before experiments.

For all experiments, cells were seeded into collagen-I coated (50 µg/mL for 1 h at 37 °C) glass bottom 24 well plates or 35 mm glass bottom dish coated with different PDMS substrates overnight at a density of 5000 cells/cm^2^. Both pHred and pHrodo labeled cells were allowed to settle for at least 15 min in the confocol microscope with 5% CO2 at 37 °C before imaging. Images were processed through the pipeline as described above, with cell masking, tracking, and background subtraction. For pHred, the intensity ratio at 561 over 405 nm (*I*_561_*/I*_405_) was calculated. For pHrodo, the average intensity at 561 nm was calculated. To obtain the absolute pH, an intracellular pH calibration buffer Kit (Invitrogen, P35379) was used with pHred or pHrodo Red AM, following the manufacturer’s protocol. Briefly, pHred cells or pHrodo dyed cells were loaded with the calibration buffer containing 10 µM valinomycin and 10 µM of nigericin at pH values of 5.5, 6.7, and 7.5 for at least 5 min. A pH calibration curve was then generated by linearly fitting the results from the calibration experiment. Single cell pHred ratio or pHrodo intensity was measured, processed, and fitted into the calibration curve to obtain single cell pH results. Relative intracellular pH change ΔpH_i_ is calculated by comparing pH_i_ before treatment or osmotic shock for each cell.

### Epi-fluorescence and confocal microscopy

For epi-fluorescence imaging, a Zeiss Axio Observer inverted, wide-field microscope using a 20x air, 0.8-NA objective equipped with an Axiocam 560 mono charged-coupled device camera was used. In some experiments, a similar microscope equipped with a Hamamatsu Flash4.0 V3 sCMOS camera was used. For confocal imaging, a Zeiss LSM 800 confocal microscope equipped with a 20x air, 0.8-NA objective, or a 63x oil-immersion, 1.2-NA objective was used. All microscopes were equipped with a CO2 Module S (Zeiss) and TempModule S (Zeiss) stage-top incubator (Pecon) that was set to 37 °C with 5% CO2 for live cell imaging. ZEN 2.6 or 3.6 Software (Zeiss) was used as the acquisition software. Customized Matlab (MathWorks) programs or ImageJ were used for image analysis subsequent to data acquisition.

### Live cell reporters, lentivirus preparation, transduction, and transfection

FUGW-pHRed (Addgene 65742) was used to measure intracellular pH. 3T3 cells stably expressing pHred were generated using lentiviral transduction and selected using flow cytometry. PIP3 indicator pcDNA3.1 AktPH-mCherry (Addgene 67301) and membrane indicator pHR GFP-CaaX(Addgene 113020) were used to generate a ratiometric image to monitor PIP3 level. Cells stably expressing GFP-CaaX were first generated using lentiviral and selected using flow cytometry. Then, GFP-CaaX cells were grown to 60-80% confluent and transfected with AktPH-mCherry using Lipofectamine 3000 reagent following the manufacturer’s recommendations.

For lentivirus production, HEK 293T/17 cells were co-transfected with psPAX2, VSVG, and the lentiviral plasmid of interest. 48 h after transfection, the lentivirus was harvested and concentrated using centrifugation. Wild-type 3T3 cells at 60-80% confluency were incubated for 24 h with 100x virus suspension and 8 µg/ml of Polybrene Transfection Reagent (Millipore Sigma).

### CRISPR knockout

3T3-Piezo1 knockout cell line was generated and validated in our previous work (*7*). In brief, single guide RNA (sgRNA) against PIEZO1 that targets the 5’-AGCATTGAAGCGTAACAGGG-PAM-3’ at Chr.8: 122513787 - 122513809 on GRCm38 was purchased from Invitrogen (A35533). The cells were transfected with the Cas9 enzyme (TrueCut Cas9 Protein v2, Thermo Fisher) and the sgRNA using Lipofectamine CRISPRMAX Transfection Reagent (Thermo Fisher) according to the manufacturer’s instructions. Transfected cells were then expanded as single-cell clones by limited dilution into 96 well plates. To evaluate the editing efficiency, GeneArt Genomic Cleavage Detection Kit (Life Technologies) was used according to the manufacturer’s instructions. Then, the following primers were used to amplify the region covering the CRISPR binding site: TATCATGGGACCTGGGCATC (forward), CAGGTGTGCACTGAAGGAAC (reverse). The knock-out of Piezo1 was confirmed by next generation sequencing (provided by Genewiz) and western blot.

### Western Blot

Western blots were performed using protocols previously described in Ref.47, 48, using NuPAGE 4-12% Bis-Tris Protein Gels (Thermo Fischer Scientific, NP0336BOX) in Invitrogen Novex Nu-Page MES SDS Running Buffer (1X, Thermo Fisher Scientific, NP0002). Primary antibody: anti-pAkt-s473 (rabbit; clone D9E; Cell Signaling; 4060; 1:1000) and anti-Akt (rabbit; clone C67E7; Cell Signaling; 4691,1:1000). Loading control: anti-GAPDH (rabbit; D16H11 Cell Signaling; 5174; 1:4000). Secondary Antibody: anti-rabbit IgG HRP-linked antibody (Cell Signaling, 7074; 1:1000) and anti-mouse IgG, HRP-linked antibody (Cell Signaling; 7076; 1:1000).

### Mathematical modeling

The three-component mechanical-chemical-ionic model was constructed as described in Eqs. 2–4. In the model, 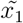 was set fixed as a control variable, and the dynamics of other two variables were solved as a system of differential equations by varying 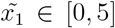, with initial conditions 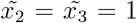 over a dimensionless timescale *τ* = 30. This timescale is sufficiently long to allow the system to reach a steady state, and the final point of the time trajectory was taken as the steady-state value. In cases without control variables, the dynamics of all three variables were solved with initial conditions 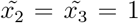. In selected tests, the effect of initial conditions was studied, and the details can be found in Supplemental Text.

### Statistical analysis

For time series plots, error bars represent the mean and the standard error of mean of at least 3 biological repeats. For scatter plots, error bars represent the mean and the standard deviation of at least 3 biological repeats. The number of cells analyzed per condition was noted in the plots or the captions. D’Agostino-Pearson omnibus normality test was used to determine whether data were normally distributed. For non-Gaussian distributions, nonparametric Mann-Whitney U-tests were used comparing two conditions, and comparisons for more than two groups were performed using Kruskal-Wallis tests followed by Dunn’s multiple-comparison test. For group comparison, two-way ANOVA was used. The statistical analysis was conducted using MATLAB 2021b (Mathworks) or GraphPad Prism 10 (GraphPad Software). Statistical significance was identified as p<0.05. ns for p > 0.05, *p<0.05, **p<0.01, ***p<0.001 and ****p<0.0001.

## Acknowledgments

This work was supported in part by NIH R01 GM134542 and R01 CA254193 (S.X.S. and K.K.). We also thank the Rockfish HPC for computational sources.

## Supplementary Figures

**Figure S1:**
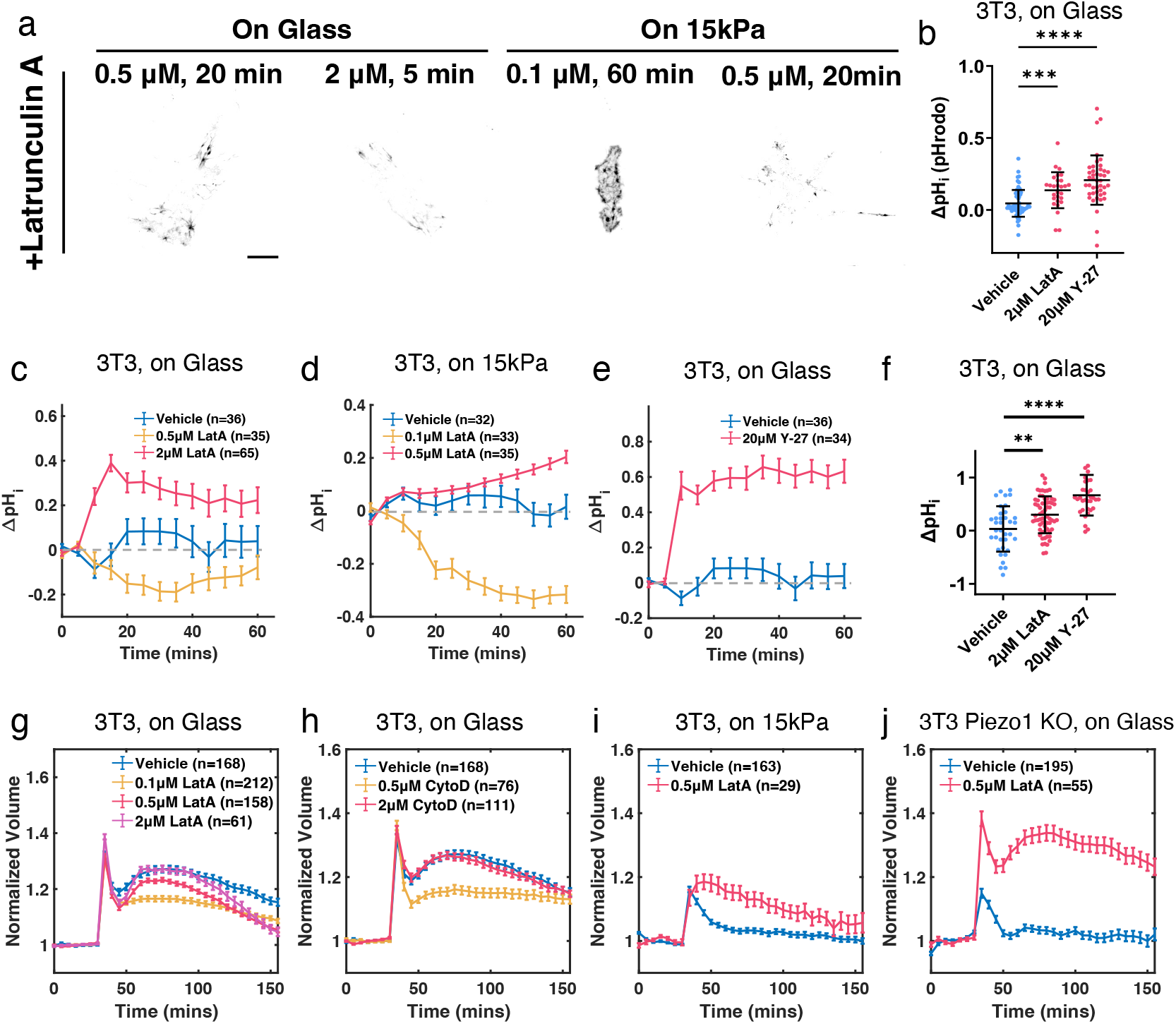
Actomyosin disruption affects intracellular pH (ΔpH_i_) and secondary volume increase (SVI) under hypotonic shock. (a) Representative confocal images of NIH 3T3 on glass stained with the live-cell F-actin probe SPY650-FastAct without treatment, and with 0.5 µM and 2 µM Latrunculin A (LatA) treatment. Cells were tracked for 1 h and the time points where most actin structures disappeared were shown. (b) ΔpH_i_ of 3T3 cells after treating with vehicle control, 20 µM Y-27632 (Y-27), and 2 µM LatA using pHrodo Red-AM. (c, d) Changes in ΔpH_i_ over time in 3T3-pHred cells treated with vehicle versus different doses of LatA on glass (c) versus on 15kPa PDMS substrates (d). (e) Changes in ΔpH_i_ over time in 3T3-pHred cells on glass treated with vehicle control and 20 µM Y-27632 (Y-27). (f) ΔpH_i_ of 3T3-pHred cells after treating with vehicle control and 20 µM Y-27 for 1 h, and 2 µM LatA for 10 min. (g, h) Volume dynamics of 3T3 treated with 0.1, 0.5, and 2 µM LatA (g), or 0.5 and 2 µM Cytochalasin D (CytoD; h). Hypotonic shock was applied at 30 min. All cells were treated 1 h prior to hypotonic shock. (i, j) Volume dynamics of 3T3 grown on 15kPa PDMS substrates (i) and 3T3 Piezo1 KO cells (j) treated 0.5 µM LatA, with hypotonic shock applied at 30 min. (b, f) Error bars represent STD. Mann Whitney U-tests. (c-e, g-j) Error bars represent the standard error of mean (SEM). (a) Scale bar = 20 µM.

**Figure S2:**
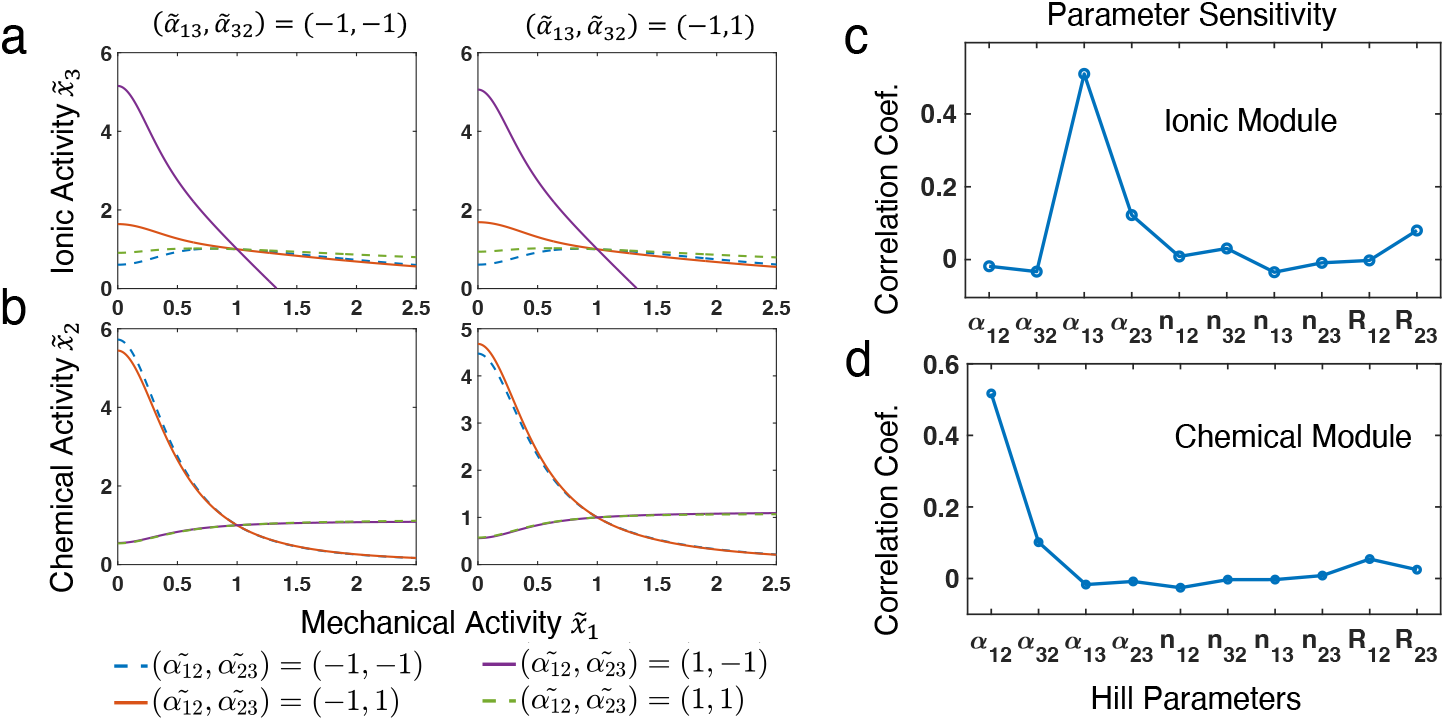
The ionic and the chemical module activity influenced by mechanical activity 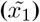 with different network structures. (a, b) The ionic (a) and the chemical module (b) activity under different network structures with actomyosin directly inhibiting NHE1 instead of activating 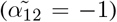. (c,d) The correlation coefficients between ionic module sensitivity (c) or chemical module sensitivity (d) and all parameters. (a-c) If not specified, the base values of all parameters are set as: 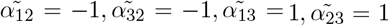, *n*_12_ = *n*_32_ = *n*_13_ = *n*_23_ = ™ 2, *R*_12_ = *R*_23_ = *R*_31_ = 1*/*5. All results are normalized to a base activity level at 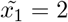.

**Figure S3:**
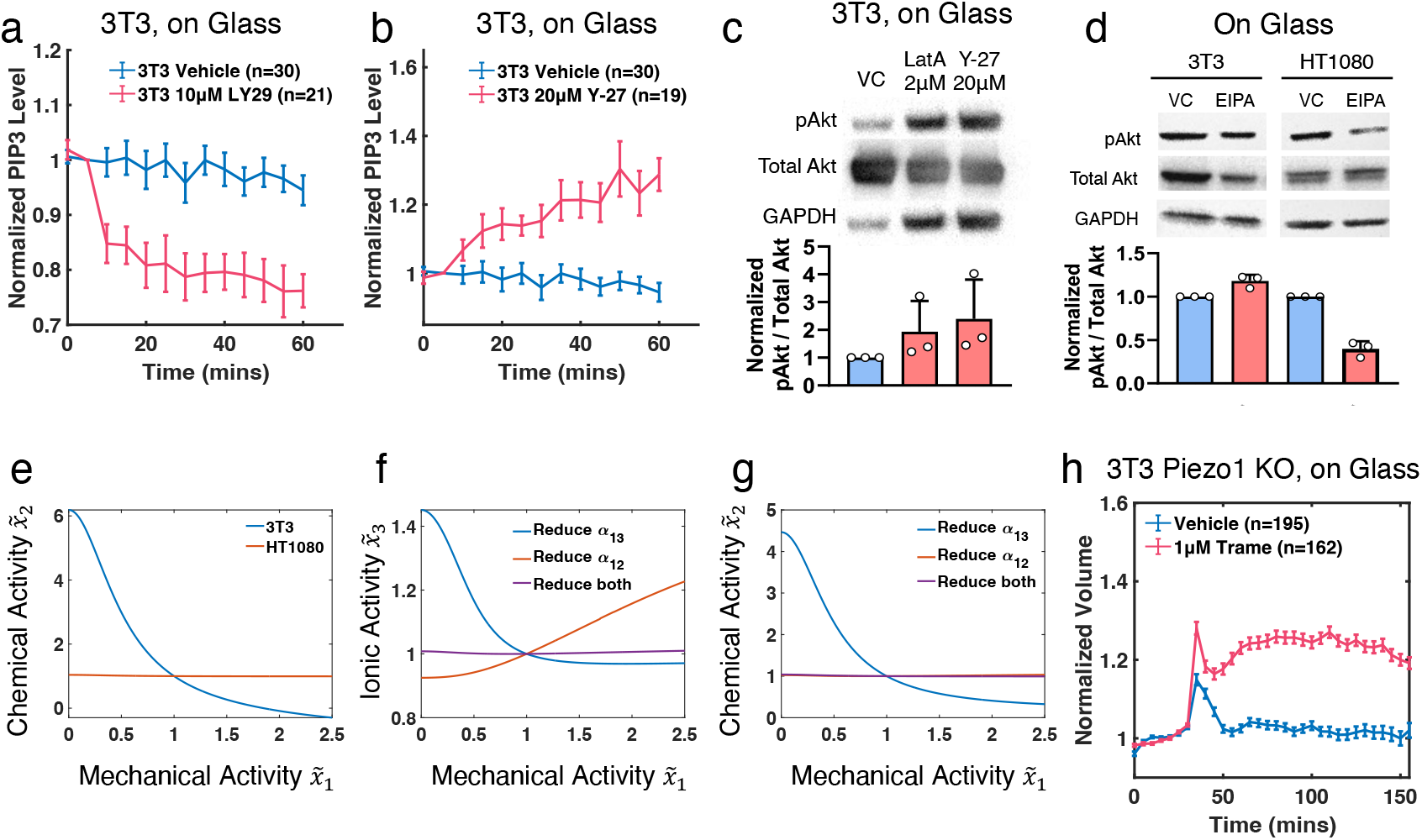
PI3K/Akt regulates chemical compensation in 3T3 but not in HT1080. (a-b) Normalized PIP3 dynamics of 3T3 cells treated with 10 µM LY29 (a) and 20 µM Y-27 (b). PIP3 level is quantified by the ratio of the PIP3 indicator (AktPH-mCherry) and the membrane indicator (GFP-CaaX) at the cell boundary. (c) Representative western blot images and quantifications of Akt activity in 3T3 cells, treated with vehicle, 2 µM LatA, and 20 µM Y-27 for 30 min. (d) Representative western blot images and quantifications of Akt activity in 3T3 and HT1080 cells treated with vehicle or 40 µM EIPA. (e) Chemical module activity of 3T3 versus HT1080 predicted by the model. (f,g) NHE1 (f) and the chemical module (g) activities modeled by reducing 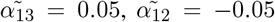, or both. Only by reducing both parameters was able to reproduce the insensitive response in HT1080. (h) Volume dynamics of 3T3 Piezo1 KO cells treated with 0.2 µM Trametinib (Trame) with hypotonic shock applied at 30 min. (e-g) 3T3 parameters are set as: 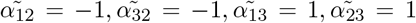, *n*_12_ = *n*_32_ = *n*_13_ = *n*_23_ = ™ 2, *R*_12_ = *R*_23_ = *R*_31_ = 1*/*5. HT1080 parameters are the same as 3T3 excepted for 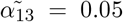 and 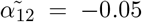. All results are normalized to a base activity level at 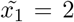. (a-c, h) Error bars represent SEM. (d) Error bars represent STD.

**Figure S4:**
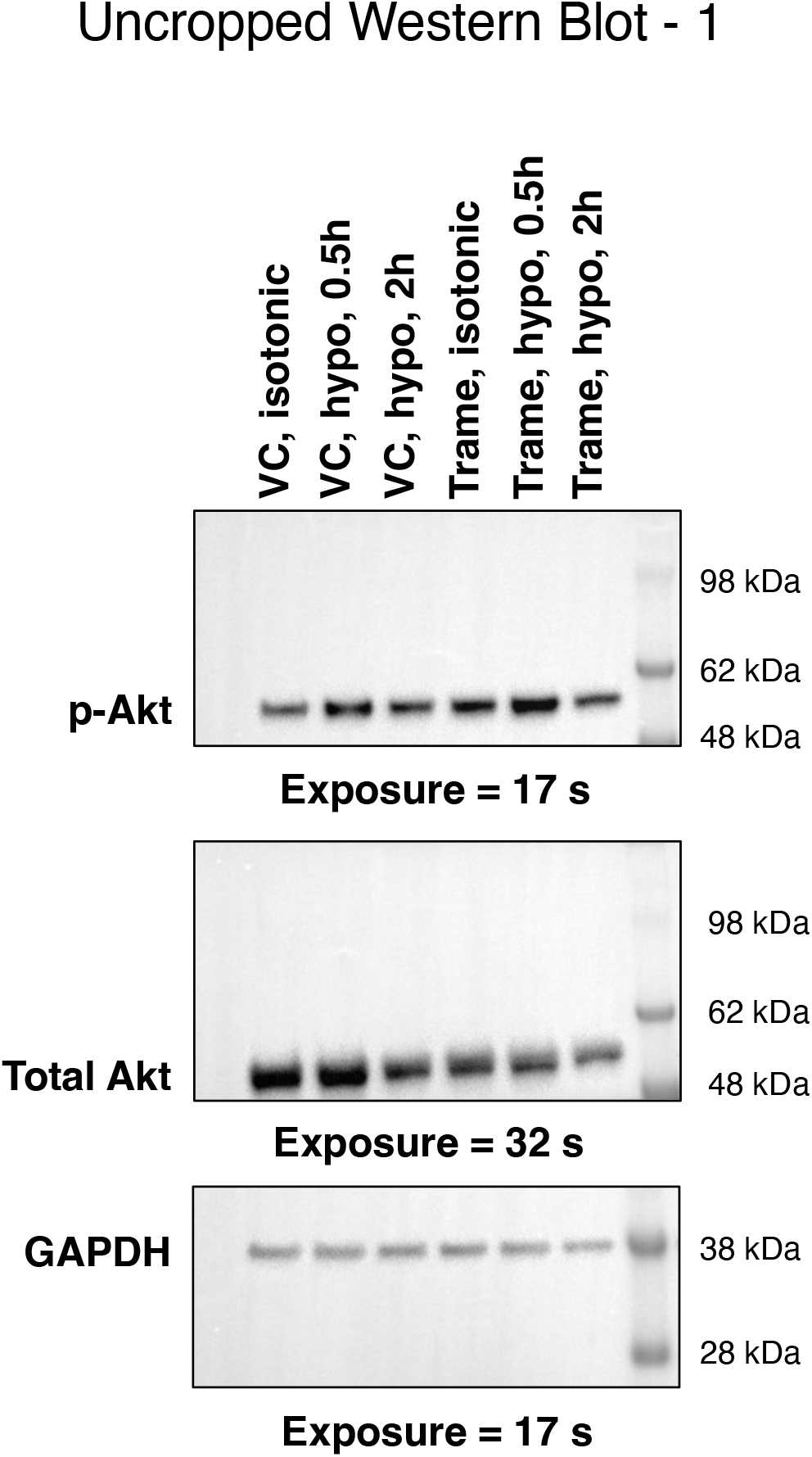
Uncropped western blots.

**Figure S5:**
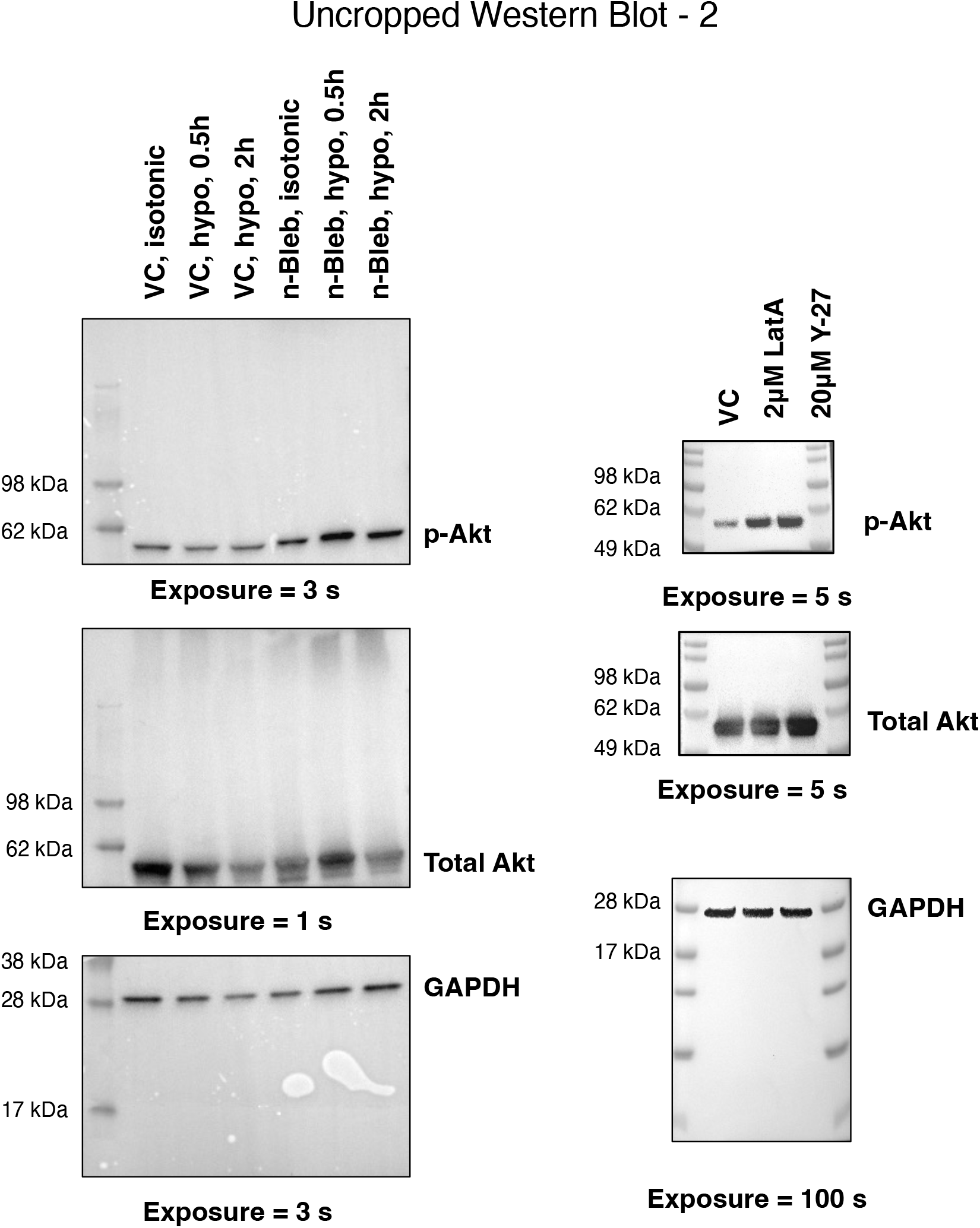
Uncropped western blots.

