## Supplementary Text for "Chemical Compensation to Mechanical Loss in Cell Mechanosensation"

### Chemical Compensation to Mechanical Loss in Cell Mechanosensation: Supplemental Text

#### Large scale unbiased model parameter sampling

The nondimensionalized 3-module network model we employed is defined as

$$\frac{k_{12}}{p_1\tau}\frac{d\tilde{x_1}}{d\tilde{t}} = 1 + \frac{\tilde{\alpha_{21}}}{1 + (R_{31}\tilde{x_2})^{n_{21}}} + \frac{\tilde{\alpha_{31}}}{1 + \tilde{x_3}^{n_{31}}} - \tilde{\beta_1}\tilde{x_1} \tag{1}$$

$$\frac{k_{23}}{p_2\tau}\frac{d\tilde{x_2}}{d\tilde{t}} = 1 + \frac{\tilde{\alpha_{32}}}{1 + (R_{12}\tilde{x_3})^{n_{32}}} + \frac{\tilde{\alpha_{12}}}{1 + \tilde{x_1}^{n_{12}}} - \tilde{\beta_2}\tilde{x_2} \tag{2}$$

$$\frac{k_{31}}{p_3\tau}\frac{d\tilde{x_3}}{d\tilde{t}} = 1 + \frac{\tilde{\alpha_{13}}}{1 + (R_{23}\tilde{x_1})^{n_{13}}} + \frac{\tilde{\alpha_{23}}}{1 + \tilde{x_2}^{n_{23}}} - \tilde{\beta_3}\tilde{x_3} \tag{3}$$

To understand globally how different parameters affect the behavior of the three-component mechano-chemical-physiological model (Fig. ST1a), we conducted a large-scale unbiased parameter search and systematically analyzed model output. A total of 50,000 parameter sets were sampled, where the parameters were selected from: $\tilde{\alpha} \in [-1.5, 1.5]$, $n \in [-2.5, -1.5]$, and $R_{12}, R_{23} \in [0.1, 10]$, all drawn from uniform distributions. These parameter ranges sufficiently cover all types of kinetics described by Hill functions in a non-dimensionalized form (*1, 2*). In this exploration, we chose the mechanical activity ($\tilde{x}_1$) as the control variable and investigated its influence on the chemical module ($\tilde{x}_2$) and the ionic module ($\tilde{x}_3$) for each parameter set. Specifically, we evaluated the steady-state values of $\tilde{x}_2$ and $\tilde{x}_3$ as functions of $\tilde{x}_1$ for a given set of parameters. Steady state solutions were taken as the end of time-dependent trajectories at dimensionless time $\tilde{t} = 30$ by solving the differential equations. Parameter sets that produced negative module activity or could not be solved with real numbers were excluded. After this filtering process, we retained 31723 parameter sets, each comprising 10 parameters, for further analysis.

Next, we defined conditions that satisfy a non-monotonic ionic (NHE1; $\tilde{x}_3$) dynamics under mechanical loss as observed in 3T3 cells. We defined these conditions as the existence of a local minimum and at least a 20% change in relative $\tilde{x}_3$ levels. A total of 760 sets of parameters satisfied these conditions. To extract potential features from these parameter sets, we applied t-distributed stochastic neighbor embedding (t-SNE) to project the high-dimensional parameter space into a lower-dimensional space for visualization (Fig. ST1b). t-SNE revealed two distinct clusters that can be easily separated, suggesting there are two parameter regimes capable of generating the non-monotonic response.

To understand which parameters drive the distinct behaviors in these two clusters, we conducted a statistical test between them. Given that the parameters were randomly sampled from a uniform distribution, we used a two-sample Kolmogorov-Smirnov test between parameters in the two cluster sets. Comparing all parameters in these two clusters revealed 4 parameters that are statistically different, including all $\alpha_{12}$, $\alpha_{13}$, $\alpha_{23}$ and $R_{23}$ (Fig. ST1c-e). Among these, $\alpha_{12}$ and $\alpha_{13}$ must be either both positive or both negative, indicating the non-monotonic response can be generated when the mechanical module inhibits or activates both of the chemical and ionic modules. However, additional conditions on $\alpha_{23}$ and $R_{23}$ must be met. In particular, $R_{23}$ is much lower in cluster 1 than cluster 2, suggesting the relative influence of mechanical module on ionic versus chemical module is a key parameter modulating the overall network dynamics.

We also examined conditions that lead to HT1080-like mechano-insensitive behavior, defined as less than a 20% change in both ionic and chemical activity across all levels of mechanical activity. In this case, we identified 1444 sets of parameters. Unlike the 3T3-type response, no visually distinguishable separation of clusters was observed after dimension reduction using t-SNE (Fig. ST2a-c). Examining the distribution of all parameters revealed that achieving an insensitive ionic and chemical response to mechanical loss requires small $\alpha_{1,2}$ regardless of their signs. While $\alpha_{1,3}$ also tends to be close to 0, many outlines at larger values can be observed (Fig. ST2c). In addition, no strong correlations between any parameters were observed in the HT1080-like response (Fig. ST2d).

### Dynamics of the three-component models

In this section, we explored the dynamical profiles of $\tilde{x}_i$ as a function of time in the three-component model. Firstly, we still make $\tilde{x}_1$ as the control variable and examine the dynamical trajectories of $\tilde{x}_2$ and $\tilde{x}_3$ at different parameter settings as discussed in the main text and shown in Figure 3b. At given $\tilde{x}_1$ and initial values at 1, $\tilde{x}_2$ and $\tilde{x}_3$ rapidly reach steady state following near monotonic trajectories (Fig. ST3a,b). Under these parameter settings, all simulations with different initial conditions generalize to the same steady state (Fig. ST3c,d), indicating the presence of a globally stable fixed point for the system at this parameter setting. From the unbiased parameter sampling,

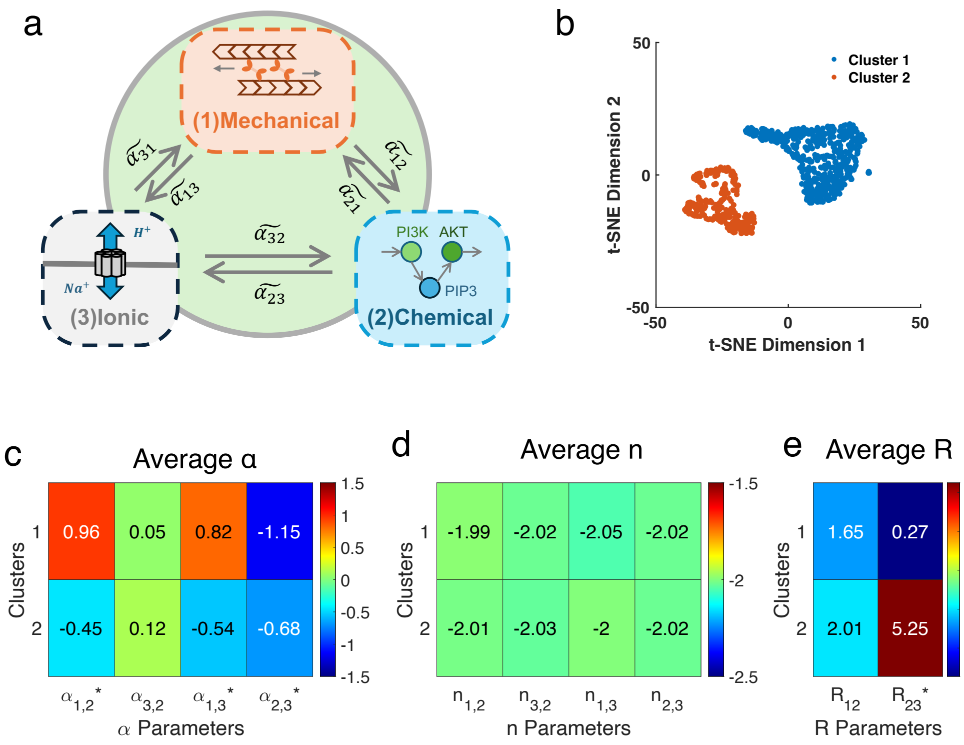


Figure ST1: **Unbiased parameter sampling reveals requirements for non-monotonic ionic dynamics.** (a) An illustration of the minimal three-component model. (b) t-distributed stochastic neighbor embedding (t-SNE) map of all parameters generating at least a 20% non-monotonic ionic activity ($\tilde{x}_3$) change as a function of mechanical activity ($\tilde{x}_1$). k-means clustering was used to identify the two clusters in the t-SNE map. (c–e) Average parameters in the two clusters. A two-sample Kolmogorov–Smirnov test was conducted between parameters in clusters 1 and 2, with * in the label indicating $p < 0.05$.

we confirmed that in most cases the initial conditions of $\tilde{x}_2$ and $\tilde{x}_3$ do not affect the steady state. In limited cases (36 out of 31723), initial values lead to distinct steady state $\tilde{x}_2$ and $\tilde{x}_3$ at certain $\tilde{x}_1$ (Fig. ST3e,f). The split of the fixed point as $\tilde{x}_1$ varies suggests the occurrence of a saddle-node bifurcation. When $\tilde{x}_1$ is fixed at $\tilde{x}_1 = 1$, the separation of steady state results show a sharp transition, suggesting cells with the same regulatory network but different concentrations or activities of the regulatory components can display distinct responses (Fig. ST3g). This might explain the emergence of phenotypes.

We next allow $\tilde{x}_i$ to freely change and explore the dynamics of all three components under different parameter sets and initial conditions. Using the same parameters as shown in Fig. ST3, no significant changes in the dynamics were observed (Fig. ST4a-c). By running a new set of unbiased parameter sampling, we found that in some cases the model generates bifurcation where the dynamics changes from stable decay to oscillatory behavior (Fig. ST4d). The generation of oscillation depends on the initial values of $\tilde{x}_i$ (Fig. ST4e). We also found that a robust limit cycle

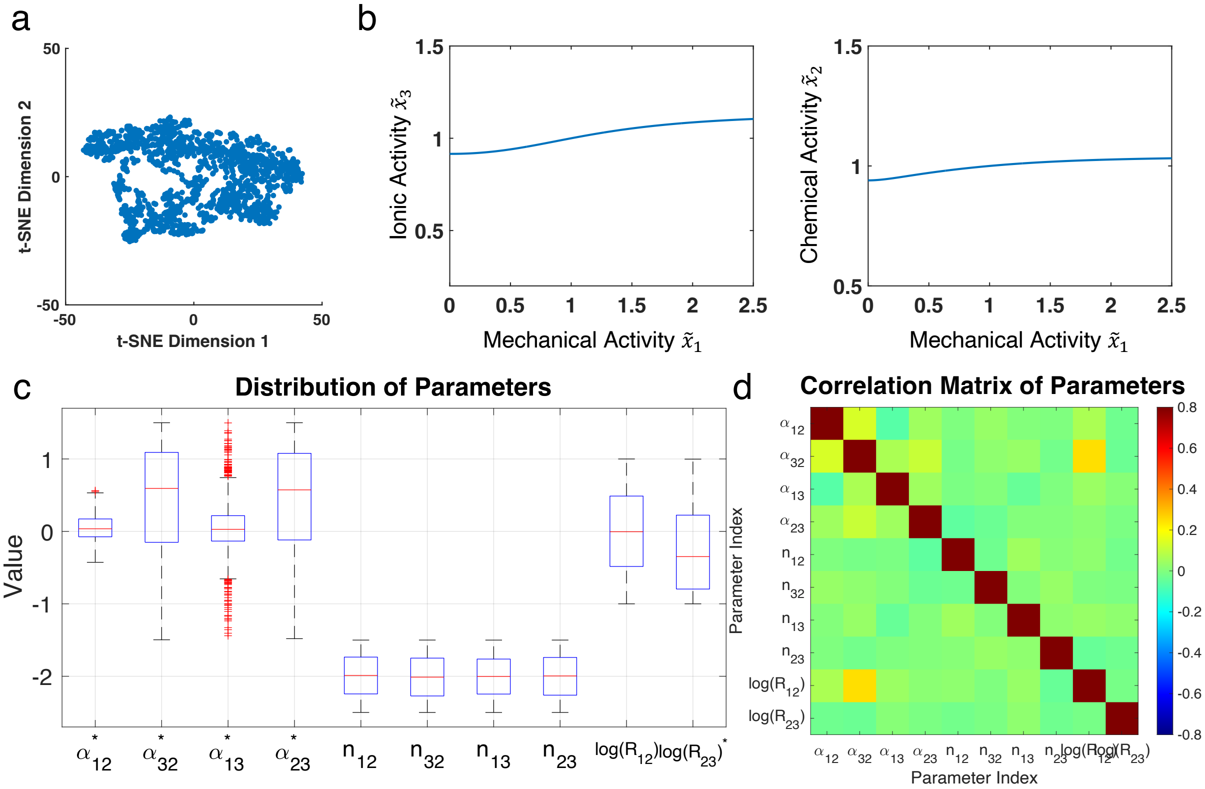


Figure ST2: **Unbiased parameter sampling reveals requirements for insensitive ionic and chemical dynamics.** (a) t-SNE map of all parameters generating an insensitive response, defined as less than a 20% change in ionic and chemical activity as functions of mechanical activity. (b) Representative ionic activity ($\tilde{x}_3$) and chemical activity ($\tilde{x}_2$) as functions of mechanical activity ($\tilde{x}_1$), using the mean parameters generating the insensitive response. All results are normalized to a base activity level at $\tilde{x}_1 = 2$. (c) Box plots of all parameters generating the insensitive response. A two-sample Kolmogorov–Smirnov test was conducted between these parameters and all sampled parameters, with * in the label indicating $p < 0.01$. (d) Correlation matrix of Pearson correlation coefficients for all parameters generating the insensitive response.

can be formed with minor variations in the parameters, and the oscillation pattern can be adjusted by varying $\tilde{\alpha}_{i,j}$ (Fig. ST4f). In sum, we conclude that our model is capable of generating different types of dynamics that observed in standard biological system.

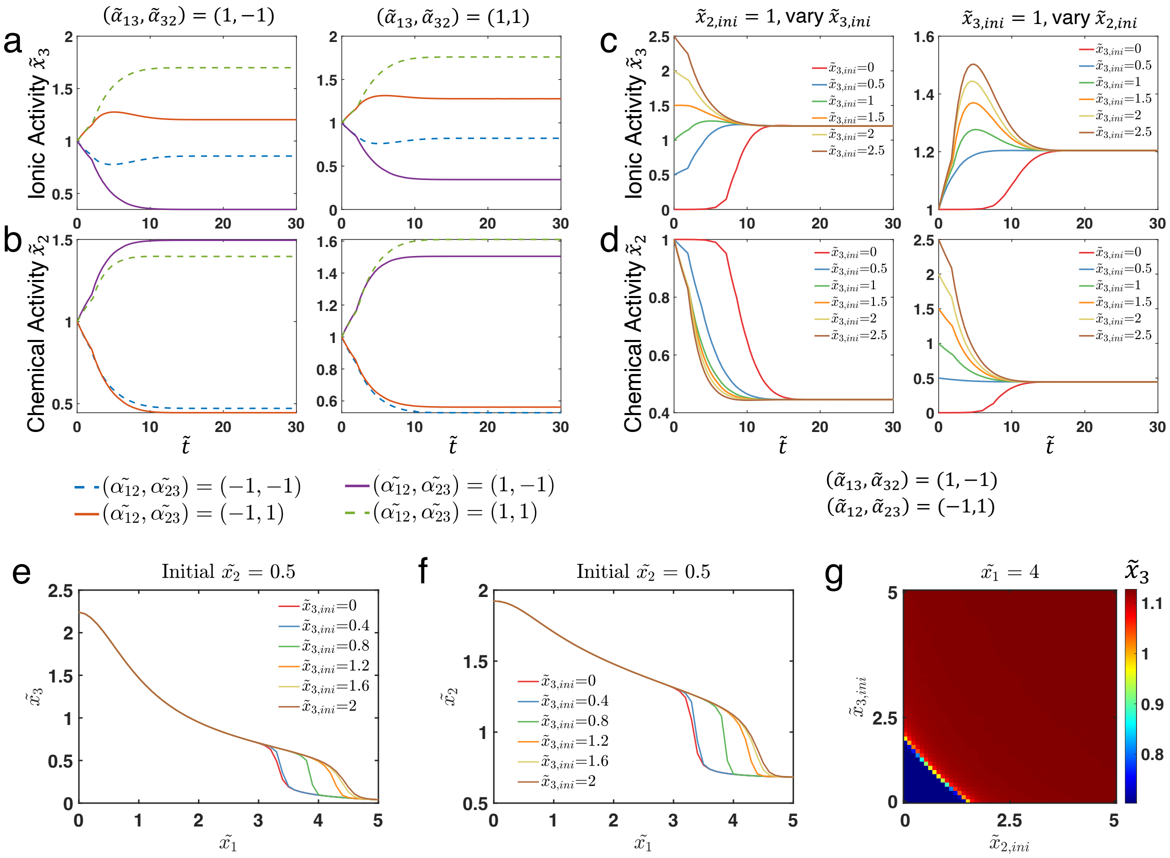


Figure ST3: **Time dependent trajectories of $\tilde{x}_2$ and $\tilde{x}_3$ and the impact of their initial conditions with $\tilde{x}_1$ as control variable.** (a, b) Time dependent trajectories of $\tilde{x}_2$ and $\tilde{x}_3$ with different parameter sets. Control variable $\tilde{x}_1 = 1$ for all conditions, and initial values $\tilde{x}_{2,ini} = \tilde{x}_{3,ini} = 1$. (c, d) The impact of initial values of $\tilde{x}_2$ and $\tilde{x}_3$ on their time dependent trajectories. Control variable $\tilde{x}_1 = 1$ for all conditions. (a-d) If not specified, the base values of all parameters are set as: $\tilde{\alpha_{12}} = -1, \tilde{\alpha_{32}} = -1, \tilde{\alpha_{13}} = 1, \tilde{\alpha_{23}} = 1, n_{12} = n_{32} = n_{13} = n_{23} = -2, R_{12} = R_{23} = R_{31} = 1/5$. (e, f, g) A representative case where initial values $\tilde{x}_{3,ini}$ affects the steady state results of $\tilde{x}_2$ and $\tilde{x}_3$. In (g), steady state values of $\tilde{x}_3$ is plotted in $\tilde{x}_{2,ini}$ and $\tilde{x}_{3,ini}$ phase space. Parameters are set as: $\tilde{\alpha_{12}} = -0.33, \tilde{\alpha_{32}} = 0.96, \tilde{\alpha_{13}} = -1.44, \tilde{\alpha_{23}} = 1.49, n_{12} = -2.1, n_{32} = -2.3, n_{13} = -1.9, n_{23} = -2.5, R_{12} = 0.38, R_{23} = 1.82$.

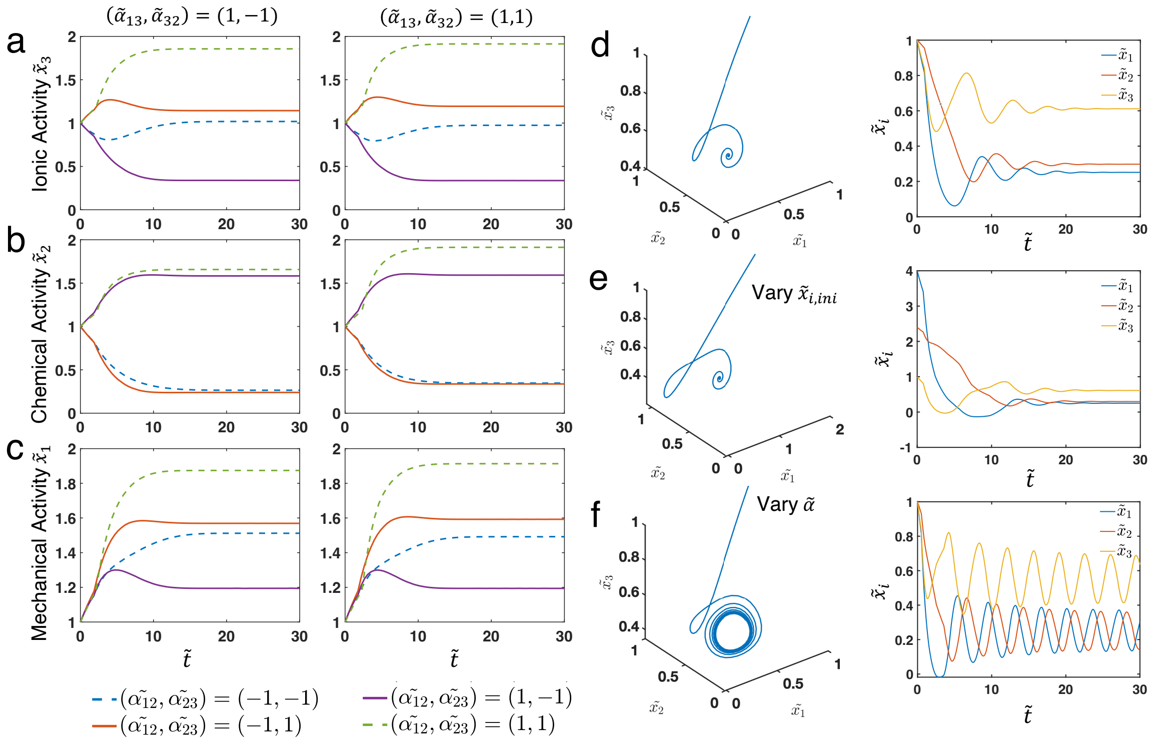


Figure ST4: **Time-dependent trajectories of $\tilde{x}_i$ without control variables and the emergence of oscillations and limit cycles.** (a-c) Time dependent trajectories of $\tilde{x}_1 - \tilde{x}_3$ with different parameter sets. Initial values $\tilde{x}_{i,ini} = 1$ for all variables. No control variables are imposed, allowing all $\tilde{x}_i$ to evolve freely. (d) Example of a parameter set that generates oscillatory dynamics in $\tilde{x}_i$. Time-dependent trajectories are shown in the $\tilde{x}_1$–$\tilde{x}_3$ phase space (left) and as separate time-series plots (right). Parameters are set as: $\widetilde{\alpha_{12}} = 1.40, \widetilde{\alpha_{21}} = -1.38, \widetilde{\alpha_{32}} = -1.10, \widetilde{\alpha_{23}} = -0.44, \widetilde{\alpha_{13}} = -1.04, \widetilde{\alpha_{31}} = -0.35, n_{12} = -2.2, n_{12} = -1.56, n_{32} = -2.43, n_{23} = -1.8, n_{13} = -2.28, n_{31} = -1.87, R_{12} = 3.13, R_{23} = 2.28, R_{31} = 2.91$. (e) Influence of initial conditions on oscillatory dynamics. Initial values are changed to $\tilde{x}_{1,ini} = 4$, $\tilde{x}_{2,ini} = 2.5$, and $\tilde{x}_{3,ini} = 1$, while all other parameters remain the same as in (d). (f) Influence of $\tilde{\alpha}_{ij}$ values on oscillatory dynamics. Initial values are $\tilde{x}_{i,ini} = 1$ for all variables. All $\tilde{\alpha}_{ij}$ values are 0.2 lower than those in (c), and all other parameters remain the same as in (c).
